# Hypercapnia Suppresses Macrophage Antiviral Activity and Increases Mortality of Influenza A Infection via Akt1

**DOI:** 10.1101/2020.02.13.946400

**Authors:** S. Marina Casalino-Matsuda, Fei Chen, Francisco J. Gonzalez-Gonzalez, Aisha Nair, Sandra Dib, Alex Yemelyanov, Khalilah Gates, G. R. Scott Budinger, Greg J. Beitel, Peter H. S. Sporn

## Abstract

Hypercapnia, elevation of the partial pressure of CO_2_ in blood and tissues, is a risk factor for mortality in patients with severe acute and chronic lung diseases. We previously showed that hypercapnia inhibits multiple macrophage and neutrophil antimicrobial functions, and that it increases the mortality of bacterial pneumonia in mice. Here, we show that normoxic hypercapnia increases viral replication, lung injury and mortality in mice infected with influenza A virus (IAV). Elevated CO_2_ increased IAV replication and inhibited antiviral gene and protein expression in macrophages *in vivo* and *in vitro*. Hypercapnia potentiated IAV-induced activation of Akt, while specific pharmacologic inhibition or shRNA knockdown of Akt1 in alveolar macrophages blocked hypercapnia’s effects on IAV growth and the macrophage antiviral response. Our findings suggest that targeting Akt1 or downstream pathways through which elevated CO_2_ signals could enhance macrophage antiviral host defense and improve clinical outcomes in hypercapnic patients with advanced lung disease.

## Introduction

Hypercapnia, elevation of the level of CO_2_ in blood and tissue, commonly occurs in patients with severe acute and chronic pulmonary disorders. It most often develops in the advanced stages of chronic obstructive pulmonary disease (COPD), the fourth leading cause of death in the U.S.(1) and third worldwide (2). Patients with COPD and other chronic pulmonary disorders are also at risk for the development of acute respiratory failure, which may be accompanied by acute or acute-on-chronic hypercapnia, often with elevations of PCO_2_ to >100 mmHg. Hypercapnia has long been recognized as a marker of poor prognosis in COPD (3-7). Importantly, COPD and the other clinical scenarios associated with hypercapnia all carry a high risk of pulmonary infection, including community-acquired pneumonia (8). Also, bacterial and viral pulmonary infections, especially influenza (9-11), are a principal cause of acute COPD exacerbations (12), and are linked to increased need for hospitalization and to mortality (13). Moreover, hypercapnia is an independent risk factor for mortality in hospitalized adults with community-acquired pneumonia (8, 14), children with adenoviral lung infections (15) and patients with cystic fibrosis awaiting lung transplantation (16).

While the association between hypercapnia and increased mortality has long been known, only recently has evidence emerged indicating that hypercapnia may play a causal role in poor clinical outcomes in patients with severe pulmonary disease and lung infection. First we found that culture of macrophages (MØs) in the presence of elevated concentrations of CO_2_ inhibited expression of tumor necrosis factor (TNF) and interleukin (IL)-6 (17), proinflammatory cytokines that are critical for host defense. The inhibitory effects of CO_2_ on TNF and IL-6 were concentration-dependent, reversible, unrelated to acidosis, and selective in that other cytokines (e.g. IL-10) were not inhibited (17). Others reported that hypercapnia similarly blocked expression of TNF and other NF-κB-dependent pro-inflammatory genes in lung epithelial cells (18). In addition, we showed that hypercapnia inhibited bacterial phagocytosis by MØs (17) and autophagy (19). Moreover, we found that hypercapnia decreased the early release of TNF, IL-6 and multiple chemokines in the lung, suppressed neutrophil function, and increased the mortality in a model of *Pseudomonas* pneumonia in mice (20). Besides effects on phagocytes, hypercapnia inhibited expression of innate immune genes, including those related to defense against bacteria, response to lipopolysaccharide, chemotaxis and cell adhesion, in human bronchial epithelial cells (21). Other recently reported effects of hypercapnia that may adversely affect clinical outcomes in patients with advanced lung disease include inhibition of Na,K-ATPase-mediated alveolar fluid clearance (22); mitochondrial dysfunction and impaired proliferation of alveolar epithelial cells and lung fibroblasts (23); and induction of skeletal muscle atrophy (24).

Influenza viruses are enveloped, negative-sense RNA viruses of the *Orthomyxoviridae* family that cause highly contagious infections of the respiratory tract (25). Influenza infections result in an estimated of 140,000–810,000 hospitalizations and cause approximately 12,000– 61,000 deaths per year (1, 26) and together with pneumonia comprise the eighth leading cause of death in the U.S. (1). Influenza viruses are the second most commonly identified cause of community-acquired pneumonia requiring hospitalization among adults in the U.S (27). Influenza infections are also among the most common causes of hospitalization for acute exacerbations of COPD (9-11, 28), which in total account for 1.9% of all hospitalizations in the U.S. and approximately 20 % of hospitalizations among individuals >65 years of age (29). The high morbidity and mortality of influenza in those with underlying chronic lung disease suggest that hypercapnia may contribute to poor clinical outcomes in such individuals. However, the impact of hypercapnia on influenza outcomes has not been examined in previously published studies. Thus, in the current investigation we explored the effects of elevated CO_2_ on influenza A virus (IAV) infection in a mouse model. Because alveolar MØs (AMØs) play a key role in limiting influenza replication and protecting against influenza-induced lung injury (30-34), and because hypercapnia suppresses MØ antibacterial function, we examined the effect of hypercapnia on the MØ antiviral response. We show that hypercapnia enhances IAV replication in the lung and increased the mortality of IAV infection in mice. Further, elevated CO_2_ increased IAV replication and inhibited antiviral gene and protein expression in MØs s *in vivo* and *in vitro*. Hypercapnic suppression of MØ antiviral activity was dependent on activation of Akt, and was blocked by pharmacologic inhibition or knockdown of Akt1. Our findings indicate that hypercapnia may play a causal role in the poor clinical outcome associated with influenza in severe lung disease, and suggest that targeting Akt1 might be a useful therapeutic strategy in patients with hypercapnia and influenza infection.

## Materials and methods

### Mice

Six- to 10-week-old C57Bl/6 mice from Jackson Laboratories were used. Experiments were performed according to a protocol approved by the Institutional Animal Care and Use Committee of Northwestern University, and according to National Institutes of Health guidelines for the use of rodents.

### Murine CO_2_ exposure

Mice were exposed to normoxic hypercapnia (10% CO_2_/21% O_2_/69% N_2_) in a BioSpherix A environmental chamber (BioSpherix, Lacona, NY). O_2_ and CO_2_ concentrations in the chamber were maintained at the indicated levels using ProOx C21 O_2_ and CO_2_ controllers (BioSpherix). Age-matched mice, simultaneously maintained in air, served as controls in all experiments. For recovery experiments, mice were exposed to 10% CO_2_ for 3 days, then infected with IAV and kept an additional day in 10% CO_2_, then returned to breathing air.

### *In vivo* influenza virus infection

C57Bl/6 mice pre-exposed to air or hypercapnia for 3 days were anesthetized with isoflurane and intubated with a 20-gauge Angiocath™ catheter. Mice were inoculated intratracheally with IAV (A/WSN/33 [H1N1]), a mouse-adapted strain, kindly provided by Robert Lamb, Ph.D., Sc.D., Northwestern University, Evanston, IL, or PBS as control were instilled as previously described (35) and returned to their previous air or hypercapnia exposure (Fig 1A). Mice were infected with 30 or 500 pfu/mouse in 50 µL of PBS.

**Fig. 1:**
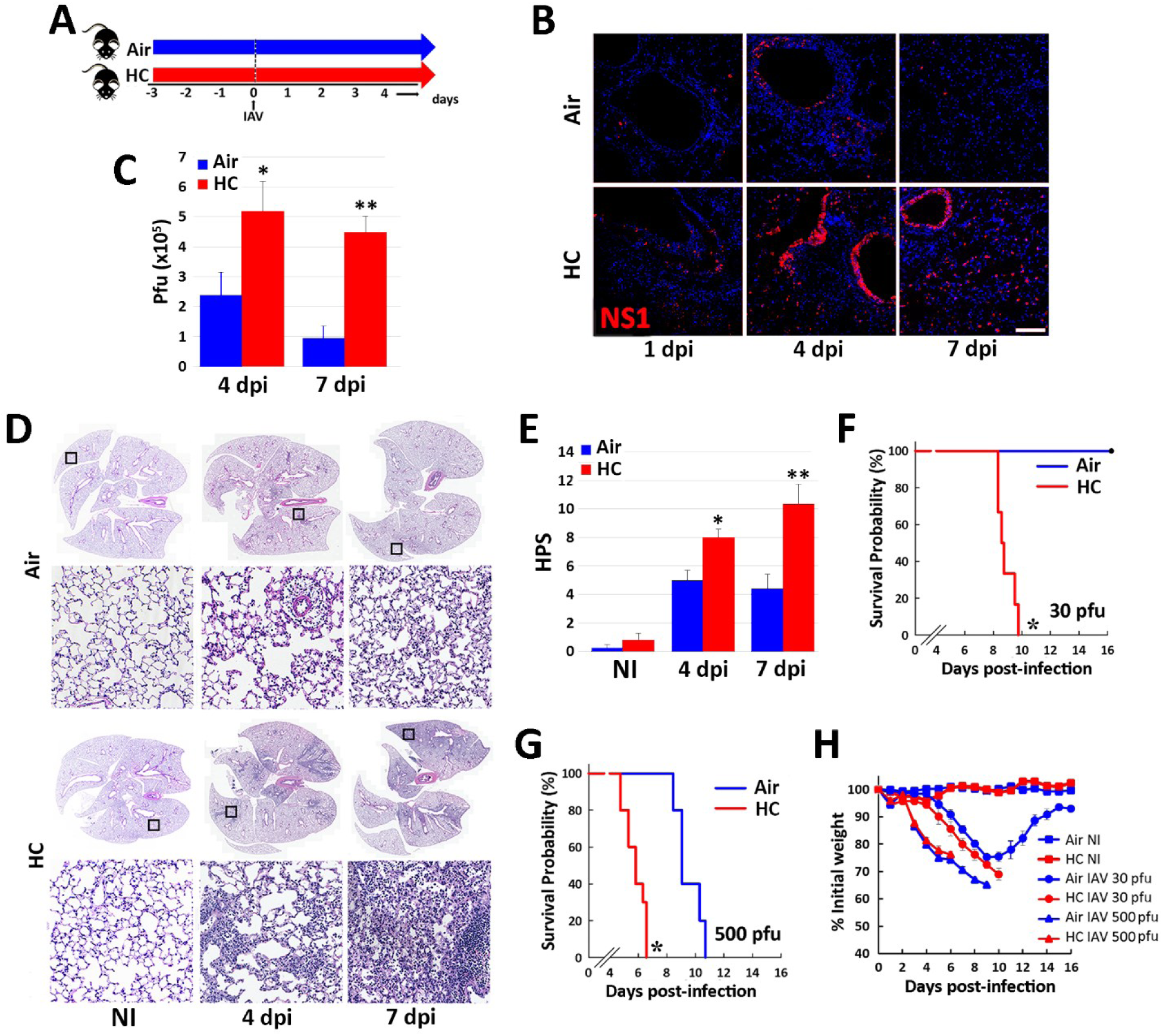
Hypercapnia increases viral protein expression, viral replication, lung inflammation and mortality in IAV-infected mice. Mice were pre-exposed to normoxic hypercapnia (10% CO_2_/21% O_2_, HC) for 3 days, or air as control, then infected intratracheally with 30 (B-F, H) or 500 (G, H) pfu IAV (A/WSN/33) per animal (A); N = 6-10 per group. Expression of viral NS1 protein (red) was assessed by immunofluorescence microscopy (IF) of lung tissue sections from mice sacrificed 1, 4 or 7 dpi; nuclei were stained with DAPI (blue) (B). Viral titers in homogenized lung tissue were determined by plaque assay at 4 and 7 dpi (C). Lungs from non-infected (NI) or IAV-infected mice harvested 4 or 7 dpi were sectioned and stained with H&E (D), and images were assessed blindly to determine histophatologic scores (HPS) for lung injury (E); *P<0.05 vs air 4 dpi, **P<0.05 vs air 7 dpi. Kaplan-Meier plots showing survival after infection with 30 (F) or 500 (G) pfu IAV; *P<0.05 vs. air by log-rank test. Body weight change over time in animals NI or infected with 30 or 500 pfu IAV (H).

### Clinical assessment of influenza A infection

Mice infected with IAV were weighed daily and monitored every 8 h for development of severe distress (slowed respiration, failure to respond to cage tapping, failure of grooming and fur ruffling). Mice that developed these signs were considered moribund and sacrificed, and the deaths were recorded as IAv-induced mortality. Mice that died between monitoring episodes were also recorded as IAV-induced mortality.

### Lung histopathology

Mice were euthanized and lungs were perfused via the right ventricle with 10 ml HBSS with Ca^2+^ and Mg^2+^. A 20-gauge angiocatheter was inserted into the trachea, and secured with a suture, the heart and lungs were removed en bloc, and lungs were inflated with up to 0.8 ml of formalin at a pressure not exceeding 16 cm H_2_O. The lungs were fixed in formalin overnight at 4°C, processed, embedded in paraffin, sectioned (4 μm thickness), and stained with H&E by the Mouse Histology and Phenotyping Laboratory at Northwestern University (Chicago, IL). Images of lungs were obtained using TissueFAXS software (TissueGnostics) at the Northwestern University Center for Advanced Microscopy (Chicago, IL). Serial images were stitched into a high-resolution macroscopic montage. Tissues were evaluated blindly using an inflammatory histopathologic score (HPS) system that assigns values of 0 to 26 (least to most severe) based on assessment of quantity and quality of peribronchial inflammatory infiltrates, luminal exudates, perivascular and parenchymal infiltrates and thickening of the membrane wall, as described previously (36, 37). This scoring system has been previously validated in other mouse models of respiratory infections (37, 38).

### Immunofluorescence microscopy in tissue sections

Sections were deparaffinized with xylene, rehydrated by using graded ethanol, and subjected to heat-induced antigen retrieval with 10 mM of sodium citrate buffer (pH 6.0). After blocking with BSA 1% (wt/vol) in PBS, tissues were incubated overnight with anti-NS1, M2, pAkt, Akt1, Akt2, or Akt3 antibodies. Then, sections were washed with PBS and Alexa–labeled secondary antibodies (1 μg/ml) were added. Co-labeling with F4/80 or YM1 (MØ markers) and SPC (alveolar epithelial cell type II marker) was achieved by the addition of anti-F4/80, YM1 or SPC antibodies respectively. Nonimmune mouse or goat IgGs were used as a negative controls. After washing, Alexa-conjugated antibodies (1 μg/ml) were added, and sections were incubated for 1 h at room temperature. Complete information in antibodies is provided in Supplementary Table 1. Nuclei were visualized with 4′,6-diamidine-2-phenylindole (DAPI) and slides were mounted with Gel/Mount (Biomeda, Foster City, CA). Fluorescent images were obtained using a fluorescence microscope Axiovert 200M (Carl Zeiss Meditec, Inc., Thornwood, NY).

### Preparation of lung homogenates for viral plaque assay and western blot

Inferior vena cava was cut and the right ventricle was perfused in situ with >1 ml of sterile PBS. Lungs were removed and kept on ice prior to and during homogenization (Tissue Tearor, 30 s) in PBS. The homogenate was split into two aliquots and an additional 1 mL of PBS was added to one of them. The cell homogenate was centrifuged (4°C, 2000 rpm for 10 min). MDCK cells were grown in 6-well plates to 100% confluence, then incubated with serial 10-fold dilutions of lung homogenate in DMEM and 1% bovine serum albumin (BSA) for 1 h (37°C). Supernatants were then aspirated, the cells were washed with PBS, 3 ml of replacement media [2.4% Avicel (IMCD, Harrington Park, NJ), 2× DMEM, and 1.5 µg of N-acetyl trypsin] were added to each well, and the plates were incubated for 3 days. The overlay was then removed and viral plaques were visualized using naphthalene black dye solution (0.1% naphthalene black, 6% glacial acetic acid, 1.36% anhydrous sodium acetate) (39). The second aliquot of lung homogenate was mixed with 0.5 mL RIPA buffer and used for immunoblot.

### Cells

For bronchoalveolar lavage (BAL) to obtain AMØs, mice were anesthetized with ketamine and xylazine. Tracheotomy was performed and a 26-gauge catheter was inserted into the trachea and secured with vinyl suture. One milliliter of ice-cold PBS was instilled and withdrawn serially three times. BAL fluid was centrifuged and AMØs, purified to ≥98% by adherence to plastic and removal of nonadherent cells, were cultured in RPMI 1640, supplemented with 10% FBS, 2 mM l-glutamine, 1 mM sodium pyruvate, 20 μM 2-ME, 100 U/ml penicillin, and 100 μg/ml streptomycin (RPMI media) (17), then rested for 24 h to avoid the transient proinflammatory profile of freshly isolated AMØs (40). BAL cells were also centrifuged in a cytospin (1000 rpm for 5 min) then fixed with 4% PFA. Human AMØs were obtained by bronchoalveolar lavage from the contralateral lung of subjects undergoing bronchoscopy for clinical diagnosis of noninfectious focal lung lesions (17, 41) under a protocol approved by the Northwestern University Institutional Review Board and cultured as for mouse AMØs.

Human monocytic leukemia THP-1 cells (American Type Culture Collection) were cultured in RPMI media and differentiated to a MØ phenotype by exposure to 5 nM PMA for 48 h (17).

### Exposure of cells to normocapnia and hypercapnia

Normocapnia consisted of standard incubator atmosphere: humidified 5% CO_2_ (PCO_2_ 36 mmHg)/95% air, at 37°C. Hypercapnia consisted of 15% CO_2_ (PCO_2_ 108 mmHg)/21% O_2_/64% N_2_. In selected experiments, Tris base was added to RPMI 1640 and HBSS so that pH was maintained at 7.4 in 15% CO_2_. Cells were exposed to hypercapnia in an environmental chamber (C-174, BioSpherix) contained within the same incubator where control cultures were simultaneously exposed to normocapnia. In all cases, cells were exposed to hypercapnia or maintained in normocapnia as control for 2 h prior IAV or IFNβ treatment. The pH of the culture media was measured with a pHOx Plus blood gas analyzer (Nova Biomedical). All media were presaturated with 5% or 15% CO_2_ before addition to the cells.

### *In vitro* influenza virus infection and IFNβ treatment

Macrophages were infected with IAV A/WSN/1933(H1N1), A/Udorn/307/1972(H2N3); or A/PR8/Puerto Rico/8/1934 (H1N1) at 1, 2, and 0.1 MOI per cell respectively, using a single-cycle protocol in RPMI media for 1 h. During the incubation at 37° C in a 5 % or 15% CO_2_ atmosphere, plates were rocked every 15 min to keep the monolayer moist and to distribute viruses evenly. Thereafter, the inoculum was removed, cells were washed twice with PBS, and fresh RPMI media was added to the plates. In other experiments, MØs in normocapnia or hypercapnia were incubated with recombinant human IFNβ (10 U/ml, R&D) for 18 h.

### Immunofluorescence microscopy in cell cultures

Cells were fixed with 4% PFA, permeabilized with 0.1% Triton X100 for 5 min, blocked with BSA 1% (wt/vol) in PBS and incubated overnight with specific antibodies against NP (1:500), NS1, M2, pTBK, pAkt, Akt1, Akt2, and Akt3. Then, sections were washed with PBS and Alexa 555–labeled anti-rabbit IgG, Alexa 488-labeled anti-mouse or Alexa 647-labeled anti-goat secondary antibodies were added. Complete antibody information is provided in Supplementary Table 1. Nuclei were visualized with DAPI. Fluorescent images were obtained using a fluorescence microscope Axiovert 200M.

### Immunoblotting

The presence of indicated proteins in lung or cell homogenates were assessed by immunoblotting using the following Abs: anti-NS1, anti-STAT1, anti-pSTAT1, and anti-Akt3; anti-RIG-I, anti-MDA5, anti-TRAF3, anti-p^S473^Akt, anti-pAkt^Y380^, pan-Akt, anti-Mx1 and anti-Akt1; anti-OAS1 and anti-viperin and anti Akt2. Complete antibody information is provided in Supplementary Table 1. Signals were detected following incubation with IRDye (1:10,000, LI-COR) Biosciences or HRP-conjugated (1:5000) secondary Abs for 1 h at room temperature using the LI-COR Odyssey Fc Imaging System. Membranes were developed and densitometry was performed using ImageStudio™ software (LI-COR).

### Quantitative real-time PCR

RNA was extracted using an RNeasy mini kit (Qiagen) and reverse transcribed to cDNA using an iScript cDNA synthesis Kit (Bio-Rad). PCR amplification was performed using CFX Connect real-time system (Bio-Rad) and the TaqMan (Applied Biosystems) or PrimeTime^®^ Predesigned (IDT) gene expression assays with FAM-labeled probes. The following primer/probe sets were used: a) TaqMan: RIG-I (Hs00204833_m1), IFNβ (Hs01077958_s1), Mx1 (Hs00895608_m1) and; b) PrimeTime: MDA5 (Hs.PT.58.1224165), IFNAR1 (Hs.PT.58.20048943), IFNAR2 (Hs.PT.58.1621113), OAS1 (Hs.PT.58.2338899), and viperin (Hs.PT.58.713843). Samples were normalized using the housekeeping gene GAPDH (Hs99999905_m1). Relative expression was calculated by the comparative CT method (ΔΔCT) (42).

### Viral adhesion and internalization

Viral adhesion and internalization were determined following pre-stablished methods (43). Briefly, to monitor viral adhesion, IAV (20 MOI) was bound on ice for 90 min to cells pre-exposed to normocapnia or hypercapnia. The cells were washed, fixed, and processed as described below for NP immunofluorescence microscopy (IF). To measure virus internalization, cells pre-exposed to normocapnia or hypercapnia were incubated IAV on ice for 90 min and either fixed immediately or shifted to 37°C for 20 min. The cells were then washed with 0.1 M glycine-0.1 M NaCl, pH 3.0, buffer for 2 min, fixed, and permeabilized. IAV localization was assessed using anti-nucleoprotein (NP) antibodies. Cells were imaged using IF, and fluorescence intensity was quantified using National Institutes of Health ImageJ software. These data are presented as corrected total cell fluorescence (CTCF), the integrated density after subtraction of background fluorescence.

### *In vitro* PI3K/Akt inhibition

THP-1 MØs were treated for 1 h with inhibitors of PI3K (LY294002, 10 µM), pan-Akt (MK2206, 5 µM), Akt1 (A674563, 50 nM) or Akt2 (CCT128930, 1 µM) then exposed to NC or HC for 2 h. Cells were then incubated in the presence or absence of IAV for 30 min or 18 h, and samples were collected for analysis of mRNA and protein.

### *In vivo* MK2206 treatment

MK2206 (MedChemExpress LLC) was administrated by oral gavage at 120 mg/kg body weight one day before hypercapnia exposure and then one day before IAV infection. Captisol (MedChemExpress LLC) was used as a vehicle for the drug and control animals were treated with vehicle only.

### Lentivirus Instillation

To knock down mouse Akt1 protein *in vivo* in lung, we generated the VSVG pseudotyped lentiviruses (109–1010 TU/ml) expressing mouse Akt1 shRNA and non-silencing shRNA as control (provided by DNA/RNA Delivery Core, Skin Biology and Diseases Resource-based Center, Northwestern University, Chicago, IL, USA). For lentivirus packaging, 293T packaging cells (Gene Hunter Corporation) were transiently transfected using Transit-2020 reagent (Mirus) with the following vectors: second generation packaging vectors psPAX2 and pMD2.G (Addgene) and third generation lentiviral expression vector pLKO (Sigma). The pLKO vectors either encoded two specific shRNAs against mouse Akt1 (Cat# TRCN0000304683) or a non-silencing control shRNA sequence (Cat# SHC002) (all from Sigma). Akt1 shRNA and control non-silencing shRNA viruses were intratracheally instilled in mice in a volume of 50 µl. Mice were exposed to 10% CO_2_, or air as control, and infected with IAV 2 weeks after lentivirus instillation. Akt1 silencing was confirmed by western blot and IF analysis, as described above.

### Statistical analysis

Data are presented as means ± SE. Differences between two groups were assessed using a Student t test. Differences between multiple groups were assessed by ANOVA followed by the Tukey–Kramer honestly significant difference test. Levene’s test was used to analyze the homogeneity of variances. Significance was accepted at p < 0.05.

## Results

### Hypercapnia increases viral growth, lung inflammation and mortality in IAV-infected mice

To investigate the effects of hypercapnia on IAV infection *in vivo*, we exposed mice to normoxic hypercapnia (10% CO_2_/21% O_2_) for 3 days prior to virus inoculation. We previously showed that exposure of mice to 10 % CO_2_ for 3 days increases arterial PCO_2_ to ∼75 mm Hg, as compared to ∼40 mm Hg in air-breathing animals, and allows for maximal renal compensation of respiratory acidosis, resulting in an arterial pH of ∼7.3 (20). Mice were then infected with IAV (A/WSN/1933) at either a low (30 pfu) or high (500 pfu) inoculum, and monitored until sacrifice at 4 or 7 days, or followed for determination of mortality due to infection (Fig. 1A). Following infection with 30 pfu IAV, NS1, a virulence factor synthesized during viral replication(44), was expressed in many more lung cells of mice breathing 10% CO_2_, as compared to air, at 1, 4 and 7 dpi (Fig 1B). By 7 dpi, NS1 expression had largely resolved in air breathing mice, but persisted in hypercapnic mice. Similarly, viral titers in homogenized lung tissue were two-fold higher at 4 dpi and remained elevated at 7dpi in hypercapnic mice, as compared to air breathing controls, in which titers declined by almost two-thirds at 7 dpi (Fig 1C). IAV-induced inflammatory lung injury was also greater in hypercapnic mice than in air breathing mice, as shown in representative images (Fig. 1D) and by histopathologic score (Fig. 1E). While air breathing mice infected with IAV at 30 pfu all survived, those exposed to 10% CO_2_ exhibited 100% mortality by 10 dpi (Fig. 1F). When infected with 500 pfu IAV, both air and 10% CO_2_ breathing mice experienced 100% mortality, but the hypercapnic mice died 3.5 to 4 days earlier (Fig. 1G). Interestingly, after IAV infection at both the low and high dose, mice lost weight at the same rate in air and 10% CO_2_ (Fig. 1H), indicating that the CO_2_ effect on mortality was not due to lesser intake of food and water in the hypercapnic mice.

Of note, when mice exposed to 10% CO_2_ were returned to breathing air one day after infection with 30 pfu IAV (Supplementary Fig S1A), NS1 expression in lung cells (Supplementary Fig S1B) and inflammatory lung injury (Supplementary Fig S1C) were reduced in comparison to mice continuously exposed to 10% after infection. Moreover, like air breathing mice, all of the animals initially exposed to 10% CO_2_ and returned to air recovered from the infection, whereas 100% mortality was again seen in those continuously exposed to 10% CO_2_ (Supplementary Fig S1D). Thus, the adverse effects of hypercapnia on viral growth and the ability of mice to survive IAV infection are reversible.

### Hypercapnia increases viral protein expression and viral replication in alveolar macrophages following IAV infection *in vivo* and *in vitro*

Examination of immunostained lung sections from IAV-infected mice exposed to air or 10% CO_2_ and sacrificed 4 dpi revealed that NS1 expression co-localized with both surfactant protein C (SPC)-positive alveolar type 2 (AT2) cells and with F4/80-positive AMØs (Fig. 2A). Interestingly, while NS1 expression was greater in both cell types in 10% CO_2_ as compared to air breathing mice, the increase was particularly notable in AMØs in the hypercapnic animals. In addition to NS1, AMØs in the hypercapnic mice exhibited increased expression of M2, a virus-encoded, multi-functional, proton-selective ion channel (39) (Supplementary Fig. S2A). Expression of NS1 and M2 was also increased in AMØs isolated by bronchoalveolar lavage (BAL) from IAV-infected hypercapnic mice at 1 dpi (Fig. 2B), and recovery of viable virus from these cells was three-fold greater than from AMØs obtained from air breathing mice (Fig. 2C). To further evaluate the effect of elevated CO_2_ on IAV growth in MØs, we infected PMA-differentiated THP-1 cells with IAV (A/WSN/1933) under normocapnic (5% CO_2_/95% air) or hypercapnic (15% CO_2_/21% O_2_/64% N_2_) conditions. After 18 h, nearly twice as many THP-1 MØs cultured in hypercapnia stained positively for nucleoprotein (NP), an IAV-encoded protein required for viral replication and assembly (45), as compared to cells cultured in normocapnia (Fig. 2D). NS1 expression (Fig. 2E) and viral titers (Fig. 2F) were also higher in THP-1 MØs cultured in hypercapnia, indicating that elevated CO_2_ enhances viral replication in these cells.

**Fig. 2:**
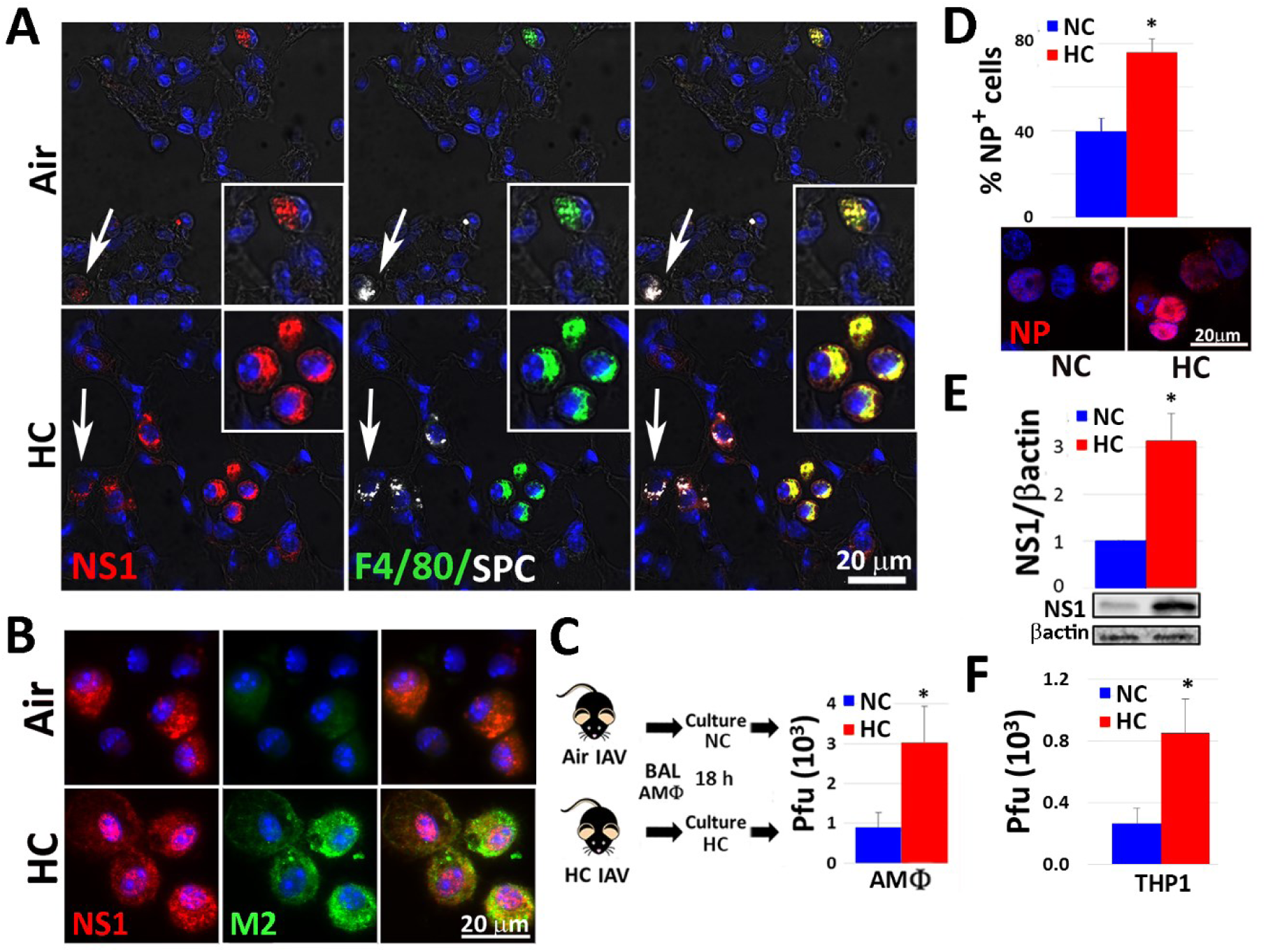
Hypercapnia increases viral protein expression and viral replication in alveolar macrophages following IAV infection of mice and in IAV-infected human THP-1 macrophages. Mice were pre-exposed to air or normoxic hypercapnia (10% CO_2_/21% O_2_, HC) and infected with IAV (30 pfu), as in Fig. 1A. Animals were sacrificed 4 dpi, and lung tissue sections were stained for viral NS1 (red), F4/80 (green, MØs), SPC (white, AT2 cells) and nuclei (blue); inserts show enlarged view of AMØs, white arrows indicate AT2 cells (A). AMØs obtained by BAL 1 dpi were stained for viral NS1 (red), M2 (green) and nuclei (blue) (B), or cultured under normocapnic (5% CO_2_/95% air, NC) or hypercapnic (15% CO_2_/21% O_2_/64% N_2_, HC) conditions for 18 h, after which viral titers in culture supernatants were determined by plaque assay (C). Differentiated THP-1 MØs were pre-exposed to NC or NC for 2 h, infected with IAV (MOI 2), and cultured in NC or HC for an additional 18 h. THP-1 MØs were then stained for viral NP (red) and the percentage of NP-positive cells was determined (D) or lysed for determination of viral NS1 expression by immunoblot (E) and viral titers in culture supernatants were determined by plaque assay (F); N = 4, *P<0.05 vs. NC + IAV.

The effect of hypercapnia was not specific to one strain of IAV, since NP expression was increased in THP-1 MØs cultured in 15% CO_2_, as compared to 5% CO_2_, and infected with A/Puerto Rico/8/1934 [H1N1] (PR8) and A/Udorn/307/1972 [H3N2] (Udorn), in addition to A/WSN/1933 [H1N1] (WSN) (Supplementary Fig. S2B).

The initial steps in replication of IAV are attachment to the surface of a target cell, followed by internalization of the virus. Thus we assayed IAV adhesion and internalization in THP-1 MØs cultured in 5% or 15% CO_2_. We found no difference in IAV adhesion or internalization, as measured by immunostaining for viral NP, at either CO_2_ concentration (Supplementary Fig. S2C-D), suggesting that hypercapnia increases viral growth in MØs by impacting processes downstream of viral entry.

### Hypercapnia inhibits the interferon pathway antiviral response to IAV in macrophages

Once IAV is internalized, viral RNA is uncoated in endosomes and binds to the cytosolic RNA helicases retinoic acid-inducible gene-I (RIG-I) and melanoma differentiation-associated protein 5 (MDA5) to initiate an antiviral signaling cascade mediated by type I interferons, interferon (IFN)-α and IFN-β. Thus, we next examined the effect of culture in 5% versus 15% CO_2_ on RIG-I and MDA5 protein expression. In the absence of viral infection, hypercapnia reduced RIG-I expression in THP-1 MØs, but did not affect the basal level of MDA5 (Supplementary Fig S3A-B). When THP-1 MØs were infected with IAV, the virus triggered marked increases in RIG-I and MDA5 mRNA and protein under normocapnic conditions, but IAV-induced gene and protein expression of both helicases was significantly inhibited in cells cultured in hypercapnia (Fig. 3A-B). Upon binding viral RNA, the helicase domain of RIG-I or MDA5 interacts with mitochondrial antiviral signaling protein (MAVS), which binds the adaptor protein, tumor necrosis factor receptor-associated factor 3 (TRAF3), resulting in phosphorylation of TANK binding kinase-1 (TBK1), activation of interferon response factor (IRF)-3 and IFR-7, and transcription of IFN-β (46). We found that hypercapnia reduced TRAF3 protein expression (Fig 3C), inhibited activation of TBK1 (Fig 3D and Supplementary Fig S3C), and reduced expression of IFN-β in IAV-infected THP-1 MØs (Fig 3E). Further, hypercapnia inhibited IAV-induced expression of IFN-stimulated genes (ISGs), including the antiviral effectors, MX dynamin-like GTPase 1 (Mx1), 2’-5’-oligoadenylate synthetase 1 (OAS1) and viperin (Fig. 3H-J). Hypercapnia also inhibited induction of RIG-I and viperin expression in response to the IAV RNA mimic and RIG-I ligand, 3p-hpRNA (Supplementary Fig. S4A). In addition, like the effects of hypercapnia on IAV growth and mortality in mice (Supplementary Fig. S1D), hypercapnic inhibition of IAV-induced viperin expression was reversible when THP-1 MØs were switched from culture in 15% CO_2_ to 5% CO_2_ (Supplementary Fig. S4B).

**Fig. 3.**
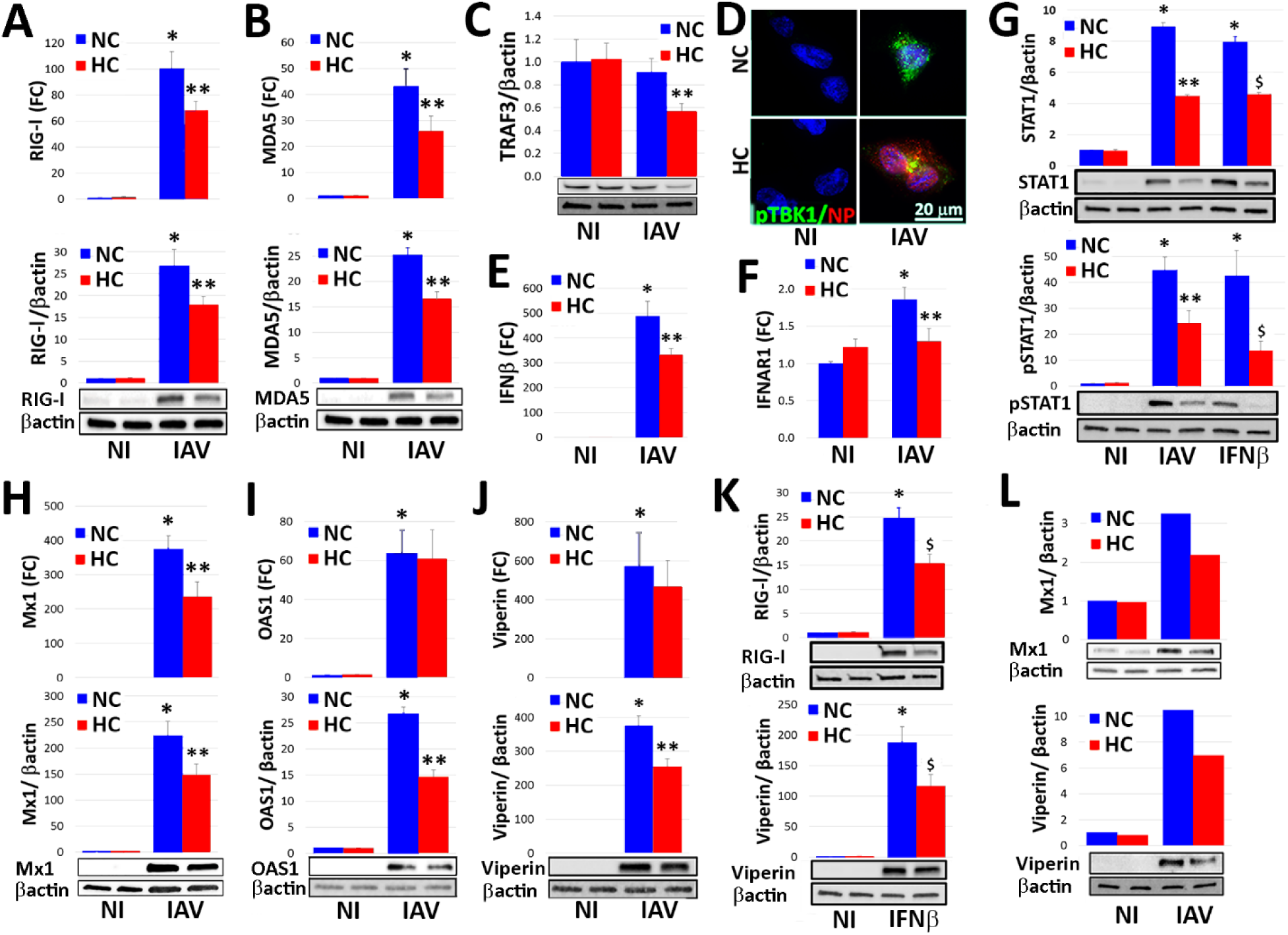
Hypercapnia inhibits IAV-induced expression and activation of key mediators of the interferon pathway antiviral response in macrophages. Differentiated THP-1 MØs or human AMØs were infected with IAV (MOI 2) (A-J, L) or stimulated with recombinant human IFN-β (10 U/ml) (G, K), and cultured in 5% CO_2_ (NC) or 15% CO_2_ (HC) for 18 h. Cells were then processed for determination of mRNA expression, expressed as fold change (FC) over non infected (NI) controls; protein expression by immunoblot, with β-actin as loading control; or IF microscopy. Expression of mRNA and/or protein is shown for RIG-I (A, K top), MDA5 (B), TRAF3 (C), IFN-β (E), IFNAR1 (F), total and phosphorylated STAT1 (G), Mx1 (H), OAS1 (I) and viperin (J, K) in control THP-1 MØs and those infected with IAV (A-C, E-J) or stimulated with IFN-β (G, K bottom); N = 5, *P<0.01 vs. NI, **P<0.05 vs. NC + IAV, $ P<0.05 vs. NC + IFN-β. Expression of viral NP (red) and phosphorylated TBK1 (pTBK1, green) was assessed by IF microscopy in control and IAV-infected THP-1 MØs (D). Expression of Mx1 and viperin protein is shown for control and IAV-infected human AMØs (L).

The foregoing results establish that hypercapnia inhibits initiation of the interferon pathway antiviral response at the level of RIG-I and MDA5, and that subsequent steps in the signaling cascade leading to antiviral effector protein expression are inhibited as well. We next assessed whether hypercapnia has additional inhibitory effects downstream of IFN-β. We found that culture of THP-1 MØs in 15% CO_2_, as compared to 5% CO_2_, reduced expression of the IFN-α/β receptor α chain (IFNAR1) (Fig. 3F), as well as both IAV- and IFN-β-induced expression and phosphorylation of STAT1 (Fig. 3G), which mediates transcription of ISGs downstream of IFN-β/IFNAR1 (47). In addition, hypercapnia inhibited expression of IFN-β-induced expression of both RIG-I and viperin in THP-1 cells (Fig. 3K). The suppressive effect of hypercapnia on IAV-induced antiviral effectors, Mx1 and viperin, was also observed in human AMØs (Fig. 3L). Thus, in addition to inhibiting initiation of the antiviral response at the level of RIG-I and MDA5, hypercapnia inhibits signaling downstream of IFN-β resulting in reduced expression of critical antiviral effectors ISGs.

### Hypercapnia’s effects on IAV replication and the macrophage antiviral response are not due to extracellular acidosis

Since raising the PCO_2_ in standard culture media lowers its pH (17), we performed experiments to determine whether the increase in viral load and inhibition of antiviral protein expression in MØs result from the reduction in pH or from the increase in PCO_2_ per se. For these experiments, media were buffered with Tris-HCl and Tris base in varying proportions to achieve pH 7.4 with 15% CO_2_, as well as pH 7.4 with 5% CO_2_ and pH 7.2 with 15% CO_2_ (the latter two conditions equivalent to the pH in non-Tris-buffered media equilibrated with 5% and 15% CO_2_-containing, respectively). When THP-1 MØs cultured in these media were infected with IAV, we observed a greater percentage of cells staining positively for NP (Supplementary Fig. S5A) and higher viral titers (Supplementary Fig. S5B) in hypercapnia (15% CO_2_) than in normocapnia (5% CO_2_), regardless of whether the pH was 7.2 or 7.4. Likewise, IAV-induced expression of RIG-I (Supplementary Fig. S5C) and viperin (Supplementary Fig. S5D) was suppressed by hypercapnia at both pH 7.2 and pH 7.4. Indeed, for each of these outcome parameters, the effect of culture in 15% CO_2_ tended to be more pronounced at pH 7.4 than pH 7.2. Thus, the effects of hypercapnia on viral growth and the antiviral response in MØs cannot be attributed to extracellular acidosis, but instead result from the higher PCO_2_, independent of pH.

### Hypercapnia enhances IAV-induced activation of Akt, which suppresses the antiviral response and increases IAV replication in MØs *in vitro* and *in vivo*

Among the intracellular signaling pathways triggered by IAV, the phosphatidylinositol 3′-kinase (PI3K)/Akt pathway is notable in that Akt activation promotes entry of IAV into cells and replication of the virus (48, 49). Thus, we investigated the impact of hypercapnia on activation of Akt in MØs *in vitro* and in the mouse lung *in vivo*. As it is unclear whether one of more of the three Akt isoforms mediates enhanced IAV cell entry and replication, we assessed Akt activation using antibodies that recognize the homologous phosphorylation sites at serine S473, S474 and S472 in the c-terminal motif and threonine T308, T309 and T305 in the T-loop of the catalytic protein kinase core in Akt1, Akt2 and Akt3, respectively (50). As expected, IAV triggered phosphorylation of Akt at S473/S474/S472 in THP-1 MØs cultured under normocapnic conditions, an effect evident 30 min and 18 h after infection (Fig. 4A and 4B). Interestingly, in the absence of infection, culture of THP-1 MØs in hypercapnia, as compared to normocapnia, also increased Akt phosphorylation at S473/S474/S472. Moreover, in IAV-infected cells hypercapnia augmented S473/S474/S472 phosphorylation, at both 30 min and 18 h (Fig. 4A and 4B). The same pattern of additive increases in S473/S474/S472 phosphorylation was seen in AMØs in lung sections from mice exposed to elevated CO_2_ and infected with IAV *in vivo* (Fig. 4C). IAV infection of MØs cultured under normocapnic conditions, and exposure of MØs to hypercapnia without infection, also triggered phosphorylation of T308/T309/T305 at 30 min, but the IAV- and hypercapnia-induced increases in Akt phosphorylation at these sites were not additive, and T308/T309/T305 phosphorylation was not sustained at 18 h in response to either IAV infection or hypercapnia (Supplementary Fig. 6A and 6B). Notably, activation of Akt by IAV and hypercapnia in THP-1 MØs occurred without significant changes in total Akt1/Akt2/Akt3 protein expression, as assessed by immunoblot using a pan-Akt antibody (Supplementary Fig. S6C).

**Fig. 4:**
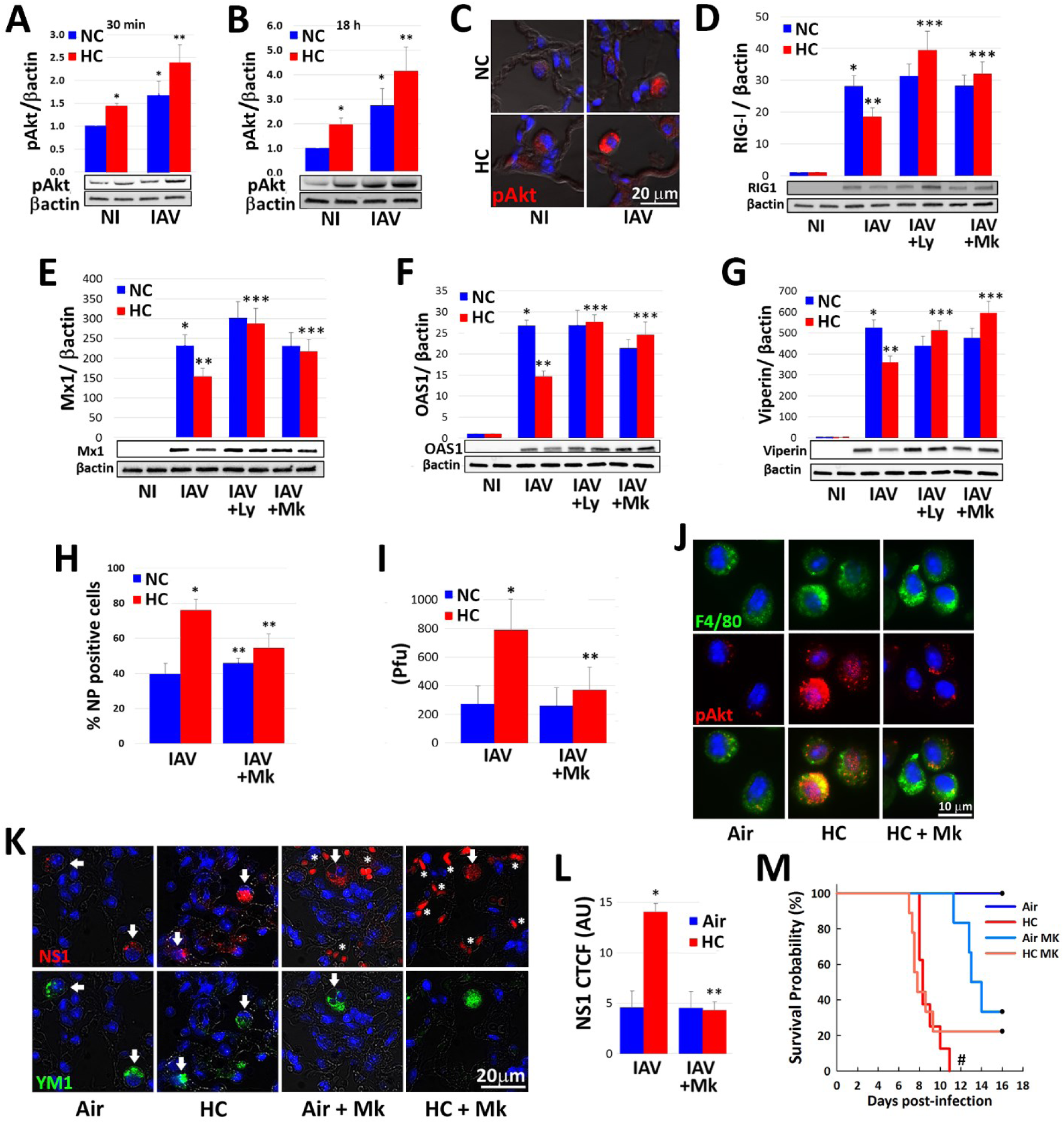
Hypercapnia potentiates IAV-induced activation of Akt, which mediates hypercapnia-induced suppression of the antiviral response and increased IAV replication in human and mouse macrophages. Differentiated THP-1 MØs were pre-exposed to 5% CO_2_ (NC) or 15% CO_2_ (HC) for 2 h, infected with IAV, and cultured for 30 min (A) or 18 h (B) prior to assessment of Ak1/Akt2/Akt3 phosphorylation at S473/S475/S472 (pAkt) by immunoblot. Additionally, mice pre-exposed to air or normoxic hypercapnia (10% CO_2_/21% O_2_, HC) and infected with IAV (30 pfu), as in Fig. 1A were sacrificed at 1 dpi, and Ak1/Akt2/Akt3 phosphorylation at S473/S475/S472 (pAkt, red) was assessed by IF microscopy in lung tissue sections; nuclei were labeled with DAPI (blue) (C). THP-1 MØs pre-exposed to NC or HC were also infected with IAV in the absence and presence of the PI3K inhibitor, LY294002 (Ly, 10 µM), or the pan-Akt inhibitor, MK2206 (Mk, 5 µM), and cultured in NC or HC for an additional 18 h, then analyzed for expression of RIG-I (D), Mx1 (E), OAS1 (F) and viperin (G) protein by immunoblot; N = 5, \**p*<0.05 vs NI; \*\**p*<0.05 vs NC + IAV; \*\*\**p*<0.05 vs HC + IAV. Cells were also immunostained for determination of the percentage of NP-positive cells by IF microscopy (H), and viral titers in culture supernatants were determined by plaque assay (I); *p<0.05 vs NC + IAV, **p<0.05 vs HC + IAV. Mice were treated with MK2206 (120 mg/kg) or vehicle control by oral gavage were exposed to HC, or air as control for 24 h, then AMØs obtained by BAL were immnostained for phosphorylation of Ak1/Akt2/Akt3 at S473/S475/S472 (pAkt, red); F4/80 (green) was used as MØ marker and nuclei were stained with DAPI (blue) (J). Other mice treated with MK2206 or vehicle were exposed to HC for 3 days, or air as control, then infected with IAV (30 pfu/animal) and maintained in air or HC for survival analysis. In addition, 5 mice in each group were sacrificed at 7 dpi, at which point viral NS1 (red) expression in lung tissue sections was assessed; YM1 was used as AMØ marker (green), nuclei were stained with DAPI (blue); white arrows indicate AMØs, asterisks denote autofluorescent RBCs (red) (K). Viral NS1 protein in AMØs in lung tissue was quantified as corrected total cell fluorescence (CTCF), expressed in arbitrary units (AU); N = 5, *p<0.05 vs NC + IAV, **p<0.05 vs HC + IAV (L). Kaplan-Meier plot showing survival after infection (M); N=8 per group), #*p*<0.05 vs Air + IAV by log-rank test.

To determine whether Akt activation plays a causal role in the hypercapnia-induced suppression of antiviral responses and the increase in viral replication, we studied cells and mice treated with MK2206, a pan-Akt1/Akt2/Akt3 inhibitor, and LY294002, an inhibitor of PI3K, the upstream kinase that activates Akt. As shown, both LY294002 and MK2206 blocked the inhibitory effect of hypercapnia on IAV-induced expression of RIG-I, Mx1, OAS1 and viperin in THP-1 MØs (Fig. 4D-4G). In IAV-infected THP-1 MØs, MK2206 also blocked the hypercapnia-induced increases in NP-positive cells (Fig. 4H) and viral titer (Fig. 4I). We also showed that when MK2206 was administered to mice by oral gavage, Akt phosphorylation at S473/S474/S472 was inhibited in AMØs obtained by BAL (Fig. 4J), and that following IAV infection the hypercapnia-induced increase in expression of NS1 was fully blocked (Fig. 4K and 4L). Treatment of mice with MK2206 also reduced the mortality of IAV infection in the setting of hypercapnia from 100% to 80%, although the change was not statistically significant (Fig. 4M). The failure of MK2206 to further improve the mortality IAV infection in hypercapnic mice probably relates to the fact that the drug itself was toxic, evidenced by the fact that treatment induced pulmonary hemorrhage and abnormalities in the liver (data not shown) and led to 70% mortality with an otherwise survivable IAV inoculum in air breathing mice (Fig. 4M).

### Akt1 mediates hypercapnic suppression of the macrophage antiviral response

The three mammalian Akt isoforms, Akt1, Akt2 and Akt3, are encoded by different genes, but share a high degree of amino acid identity (51). Although the majority of the literature does not distinguish between the three isoforms, there is a growing list of differences among them, and at least in the context of tumor initiation and differentiation, they have low functional redundancy (52, 53). Thus, we sought to determine which Akt isoform(s) are important in hypercapnic suppression of antiviral mechanisms in the MØ. First, we found that all three isoforms are expressed in mouse AMØs (Fig. 5A and Supplementary Fig. S6D) and THP-1 MØs (Supplementary Fig. S6E-G), as assessed by immunofluorescence microscopy and immunoblot using isoform-specific antibodies. Next, we showed that the Akt1-selective inhibitor, A674563, blocked hypercapnic suppression of IAV-induced RIG-I and viperin expression in THP-1 MØs, but the Akt2-selective inhibitor, CCT128930, did not (Fig. 5B-5C). The Akt1 inhibitor also blocked the increase in viral titer caused by hypercapnia, whereas the impact of the Akt2 inhibitor was less clear, since it caused an increase in viral titer under normocapnic conditions (Fig. 5D). Because selective Akt3 inhibitors are not available, we could not perform pharmacologic inhibition studies targeting Akt3. To further evaluate the role of Akt1 as a mediator of hypercapnia’s effects, we constructed a lentivirus containing an Akt1-targeted shRNA, whose effectiveness in reducing Akt1 protein expression we confirmed in cultured MLE cells (Supplementary Fig. S7A). Importantly, intranasal inoculation of mice with this lentivirus resulted in a persistent reduction in Akt1 in AMØs to less than half the level in cells from mice treated with a control lentivirus containing non-silencing shRNA (Fig. 5E and Supplementary Fig. S7B). We then cultured AMØs obtained from mice treated with lentivirus containing either the Akt1-targeted shRNA or control shRNA with IAV under normocapnic or hypercapnic conditions, and infected them with IAV. In this experiment, the hypercapnia-induced increase in NS1 expression was reduced by half in AMØs in which Akt1 had been knocked down (Fig. 5F). This confirms that Akt1 mediates the suppressive effect of hypercapnia on the MØ antiviral response. Finally, we tested whether shRNA knockdown of Akt1 in vivo would protect mice from hypercapnia-induced mortality following IAV infection. This experiment was complicated by the fact that treatment of mice with both the control and Akt1-targeted lentivirus caused inflammatory lung injury, such that both air and 10% CO_2_ breathing mice subsequently infected with IAV exhibited significant mortality, thus obscuring whether the severity of IAV infection could be reduced by Akt1 knockdown *in vivo* (data not shown).

**Fig. 5:**
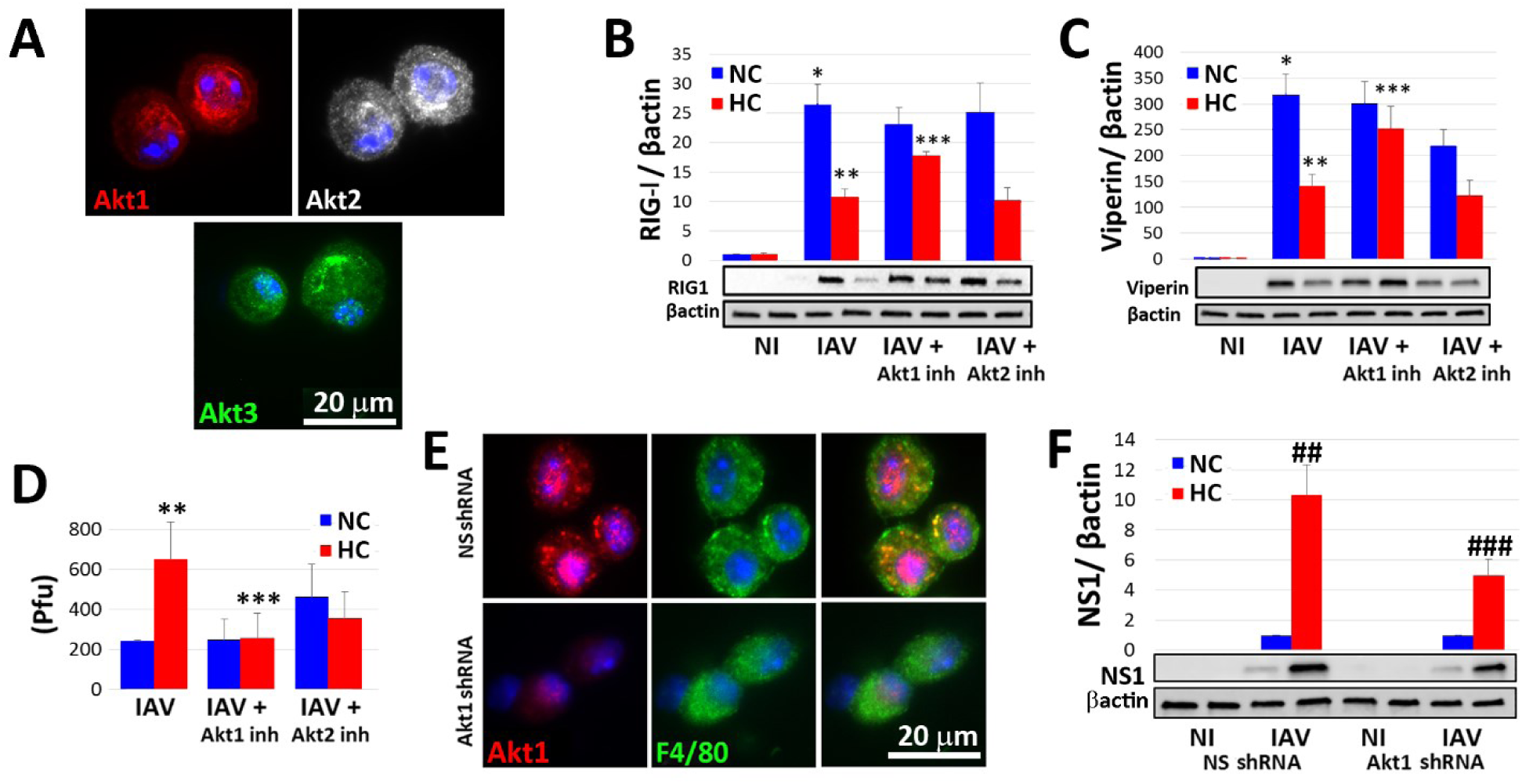
Hypercapnia-induced suppression of the macrophage antiviral response is mediated by Akt1. AMØs obtained by BAL from untreated mice were immunostained with isoform-specific antibodies for Akt1 (red), Akt2 (white) and Akt3 (green); nuclei were stained with DAPI (blue) (A). Differentiated THP-1 MØ were exposed to 5% CO_2_ (NC) or 15% CO_2_ (HC) in the absence or presence of specific inhibitors of Akt1 or Akt2, A674563 (50 nM) and CCT128930 (1 µM), respectively, and infected with IAV. Expression of RIG-I (B) and viperin (C) protein was assessed by immunoblot, and viral titers in culture supernatants were assessed by plaque assay (D); N = 4, \**p*<0.05 vs NC NI; \*\**p*<0.05 vs NC + IAV; \*\*\**p*<0.05 vs HC + IAV. In additional experiments, mice were infected intranasally with lentivirus containing non-silencing (NS) shRNA or Akt1-targeted shRNA, and 21 days later AMØs were collected by BAL and immunostained for Akt1; F4/80 (green) was used as MØ marker and nuclei were stained with DAPI (blue) (E). Alternatively, AMØs from the shRNA-treated animals were cultured in NC or HC, infected with IAV, and 18 h later processed for quantitation of viral NS1 protein by immunoblot, with β-actin as loading control (F); N = 4, ##*p*<0.05 vs NS shRNA + IAV in NC, ### *p*<0.05 vs NS shRNA + IAV in HC.

## Discussion

To our knowledge, our study is the first to examine the effect of hypercapnia on host defense against viral infection. We have shown that hypercapnia, at a level of CO_2_ relevant to patients with severe acute and chronic lung disease, increases IAV replication, virus-induced lung injury, and the mortality of IAV infection in mice. Hypercapnia increased expression of viral proteins in both lung epithelial cells and AMØs, and the increases were particularly striking in MØs. For this reason, and because AMØs play a critical role in host defense against IAV (30-34) we focused attention on the effects of elevated CO_2_ on the MØ response to IAV. We found that hypercapnia increased viral replication in mouse AMØs and differentiated human THP-1 MØs. Hypercapnia inhibited IAV-induced expression of the helicases RIG-I and MDA5, as well as downstream interferon pathway signaling, resulting in reduced MØ expression of multiple antiviral effector proteins. We further showed that hypercapnia enhances IAV-induced activation of Akt, and that the isoform Akt1 mediates most of all of the effects of elevated CO_2_ on the antiviral response and on viral growth in the MØ.

Of note, the effect of hypercapnia on antiviral activity and IAV replication in cultured MØs was unrelated to extracellular acidosis. These results are similar to our previous observations that hypercapnia suppressed IL-6 expression and autophagy in MØs in a manner unrelated to changes in extracellular (or intracellular) pH (17, 19). They also parallel our previous *in vivo* findings, in which exposure of mice to 10% CO_2_ increased the mortality of *Pseudomonas* pneumonia to an identical degree in the setting of acute and chronic respiratory acidosis (arterial pH 7.15 and 7.3, respectively). The observation that hypercapnic suppression of MØ host defense against IAV is not a function of pH suggests the possibility that antiviral signaling pathway(s) within the cell are altered by molecular CO_2_ itself.

A critical role for AMØs in host defense against influenza viruses is well-established, as depletion of AMØs using clodronate or various genetic strategies increases viral replication, lung injury and mortality following IAV infection in pigs and mice (30-34). While influenza viruses readily infect MØs, some studies show that IAV infection of AMØs is abortive, as a result of block(s) to production of live virus (54-57). In mice infected with IAV in the present study, AMØs expressed viral NP, NS1 and M2 proteins and, when isolated, released viable virus at titers orders of magnitude higher than the original inoculum, consistent with productive infection of AMØs *in vivo*. Importantly, exposure of mice to 10% CO_2_ increased NP, NS1 and M2 protein expression and titers of IAV released by AMØs in comparison to levels seen in air-breathing mice, indicating that hypercapnia inhibited pathway(s) that otherwise restrain viral replication in the AMØ.

To elucidate the mechanisms by which elevated CO_2_ inhibits MØ host defense against influenza, we used differentiated human THP-1 MØs and mouse BMDM, in both of which hypercapnia increased viral protein expression and IAV replication, like in the AMØ. The first critical step in the process of IAV Infection is binding of the virus to sialic acid, galactose-type lectins or mannose receptors in host cells (58, 59). We found that hypercapnia did not enhance adhesion of IAV to THP-1 MØs, nor did it enhance the next step, internalization of the virus, thus implicating downstream steps in the MØ response to infection. When IAV enters the cell, it rapidly induces expression of RIG-I and MDA5, which then bind viral RNA and initiate the antiviral response (60-62). Notably, exposure of THP-1 MØs to elevated CO_2_ both reduced basal expression of RIG-I and blunted the IAV-induced increase in RIG-I and MDA5 expression. Hypercapnia also reduced expression of the downstream signaling component, TRAF3, and activation of its target, TBK-1, in IAV-infected THP-1 MØs. TRAF3 and activated TBK-1 are required for activation of IRF-3 and IRF-7, which drive transcription of IFN-α and IFN-β (63, 64). Elevated CO_2_ also inhibited signaling downstream of IFN-β by reducing expression of its receptor, IFNAR1, and blunting activation of STAT-1, leading to decreased expression of ISGs, including the antiviral effectors, Mx1, viperin and OAS-1 (65-67). Notably, hypercapnia inhibited activation of STAT1 and expression of antiviral ISGs in non-virus-infected MØs stimulated with exogenous IFN-β, indicating that elevated CO_2_ blocks signaling downstream as well as upstream of IFN-β.

Among the IAV gene products upregulated in MØs by hypercapnia, NS1 is particularly important as a virulence factor, as it interferes with antiviral signaling at multiple levels (68). NS1 blocks TRIM25-mediated ubiquitinaton of RIG-I, which is required for its activation and initiation of the antiviral response (69) and also inhibits activation of IRF-3, thereby reducing transcription of IFN-α and IFN-β (70). Further downstream, NS1 inhibits IFN-β-induced phosphorylation of STAT1 and STAT2, preventing their nuclear translocation and DNA binding, and decreasing expression of STAT1/2-dependent antiviral genes (71). Another important mechanism is that NS1 inhibits the processing of cellular mRNAs by binding cleavage and polyadenylation specificity factor (CPSF 30), resulting in generally reduced expression of host genes, including IFNs and antiviral ISGs (72). Finally, at the post-translational level, NS1 inhibits the activity of the antiviral ISGs, PKR and OAS. Upon binding viral RNA within the cell, PKR phosphorylates translation initiation factor eIF2, which nonspecifically represses protein synthesis, thus inhibiting viral replication (73). OAS catalyzes the formation of 2′-5′-polyA oligomers, which activate RNAse L, a potent repressor of viral infection due to its ability to degrade single-stranded RNA (67, 74). Thus, by increasing NS1 expression, hypercapnia disrupts IFN pathway signaling and antiviral activity by multiple distinct NS1-dependent mechanisms. Additionally, hypercapnia inhibits MØ antiviral responses independently of NS1 or other viral proteins, since STAT1 activation and production of antiviral proteins triggered by IFN-β and 3p-hpRNA (in the absence of IAV infection) were decreased by elevated CO_2_ as well.

Upon infection, IAV activates the PI3K-Akt pathway in host cells in a biphasic manner. First, there is early, transient activation of the pathway associated with viral attachment and endocytosis, and beginning 2-3 h later, a second wave of sustained activation results from binding of viral NS1 to the p85 regulatory subunit of PI3K (75, 76). Notably, previous studies have not addressed which of the three Akt isoforms is activated in IAV-infected cells. In the current investigation, we showed that Akt1, Akt2 and Akt3 were all expressed in mouse AMØs and human THP-1 MØs, and confirmed PI3K-Akt activation at both the early (30 min) and later (18 h) time points after IAV infection in THP-1 MØs by showing serine phosphorylation of Akt1-S473/Akt2-S474/Akt3-S472 (with a phospho-specific antibody that does not distinguish between Akt1, Akt2 and Akt3, as used in prior studies by other groups). Notably, in the absence of infection, exposure to elevated CO_2_ also triggered phosphorylation of Akt1-S473/Akt2-S474/Akt3-S472, and moreover, hypercapnia augmented IAV-induced Akt activation following viral infection. (Akt was also phosphorylated at Akt1-T308/Akt2-T309/Akt3-T305 independently by elevated CO_2_ and IAV by 30 min after infection, but the effects of hypercapnia and IAV were not additive, and neither stimulus increased phosphorylation at this site over baseline at 18 h. Given that hypercapnia augments NS1 expression, the increased phosphorylation of Akt in IAV-infected MØs cultured in elevated CO_2_ may result from enhanced PI3K activation due to NS1-p85 binding as above, or to a direct interaction of NS1 with Akt (77).

Using a PI3K and pan-Akt pharmacologic inhibitors, we were able to block the suppressive effect of hypercapnia on IAV-induced expression of RIG-I and antiviral ISG effectors, and prevent the hypercapnia-induced increases in viral protein expression and IAV replication in THP-1 MØs. Treatment with the pan-Akt inhibitor MK2206 decreased the mortality of IAV infection in hypercapnic mice from 100% to 80%, a non-significant difference likely due to the fact that the inhibitor was itself toxic *in vivo*, causing high mortality of an otherwise nonlethal IAV infection in air-breathing animals. With the use isoform-selective inhibitors, we showed that hypercapnic suppression of antiviral host defense in THP-1 MØs was mediated by Akt1, not Akt2. Due to lack of a selective inhibitor, we were unable to evaluate the role of Akt3, although the fact that the Akt1-selective inhibitor fully blocked the hypercapnia-induced increase in IAV replication suggests that Akt3 may not play a role in mediating effects of elevated CO_2_. The importance of Akt1 as a mediator of CO_2_’s effects is further supported by the observation that Akt1- but not Akt2-targeted shRNA reversed hypercapnic suppression of antiviral protein expression and the hypercapnia-induced increase in NS1 expression in AMØs. Of note, Murray *et al* previously reported that siRNA knockdown of Akt1, but not Akt2, inhibited replication of IAV in several malignant epithelial cell lines (Hep3B, HepG2 and TZM-bl) (78). To our knowledge, in addition to showing a role for Akt1 in hypercapnic suppression of antiviral host defense, ours is the first study to specifically implicate Akt1 as the isoform critical for IAV growth in non-malignant cells and the myeloid lineage in particular.

Together with our previous finding that hypercapnia worsens the mortality of bacterial pneumonia in mice (20), our current observations on hypercapnic suppression of host defense against IAV suggest a causal role for hypercapnia in poor clinical outcomes in patients with severe COPD and other advanced lung diseases who develop pulmonary infections (3-11, 14, 16, 79). Notably, the hypercapnia-induced defects in antiviral host defense were reversible. In our previous studies, the increase in mortality of *P. aeruginosa* pneumonia in mice was similarly reversible (20), as were hypercapnia-induced inhibition of IL-6 expression (17) and bacterial-triggered autophagy (19) in MØs. The reversibility of elevated CO_2_-induced defects in antiviral and antibacterial immunity may in part explain why use of noninvasive ventilation to reduce arterial PCO_2_ prolonged the time to hospital readmission and decreased mortality in patients with severe COPD and chronic hypercapnia (80, 81).

In summary, in the first-ever study to examine the effect of hypercapnia on host defense against viral infection, we have shown that normoxic hypercapnia increases viral replication, lung injury and mortality in mice infected with IAV. We found that elevated CO_2_ increases IAV replication and inhibits antiviral gene and protein expression in MØs *in vivo* and *in vitro*. Hypercapnia potentiated IAV-induced activation of Akt, while specific pharmacologic inhibition or shRNA knockdown of Akt1 blocked hypercapnia’s effects on IAV growth and the macrophage antiviral response. Our findings suggest that targeting Akt1 or downstream pathways through which elevated CO_2_ signals may be a useful strategy to enhance MØ antiviral host defense and improve clinical outcomes in hypercapnic patients with advanced lung disease.

## Acknowledgments

Histology services were provided by the Northwestern University Mouse Histology and Phenotyping Laboratory supported by NCI P30-CA060553 awarded to the Robert H Lurie Comprehensive Cancer Center.

