## Supplementary figures and table for "Hypercapnia Suppresses Macrophage Antiviral Activity and Increases Mortality of Influenza A Infection via Akt1"

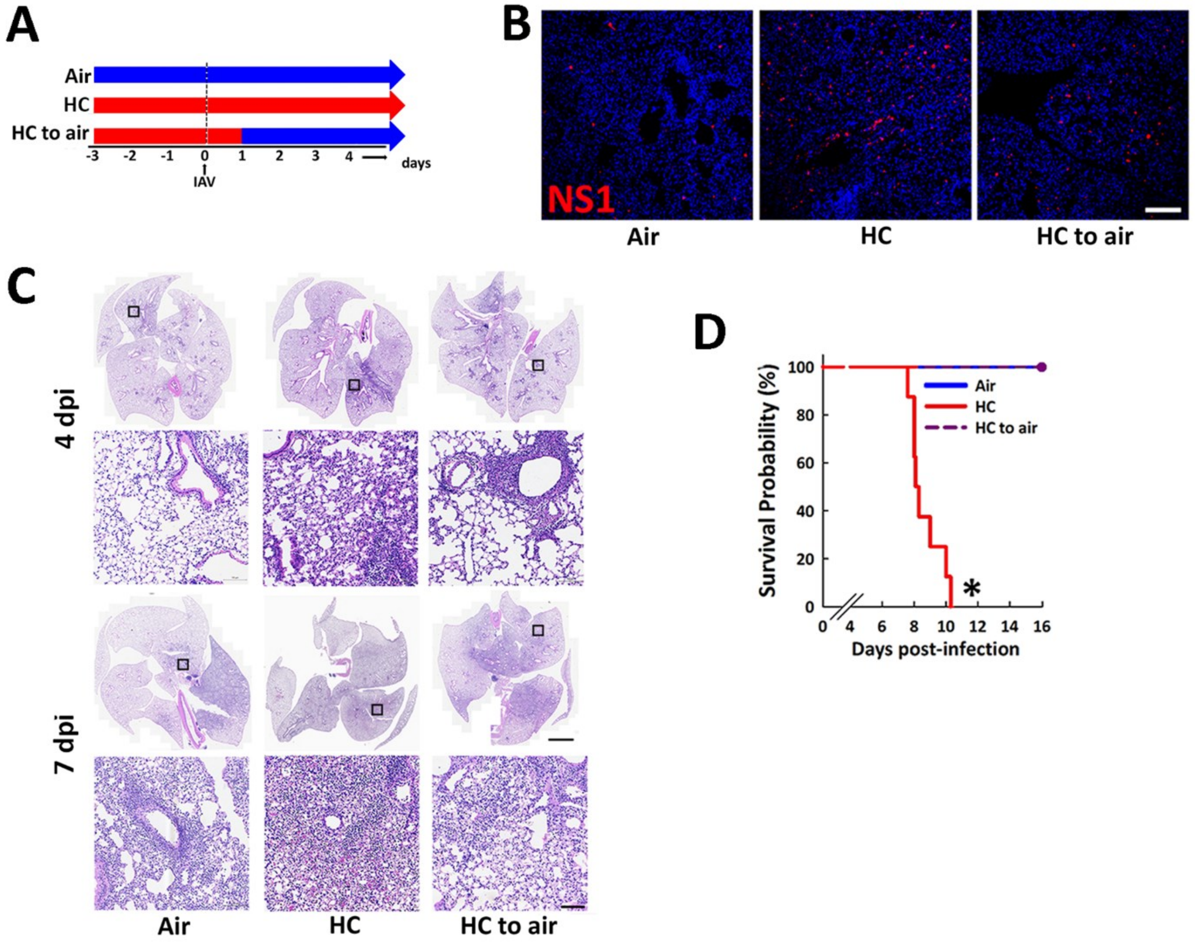


**Fig. S1:** **Hypercapnia-induced increases in viral protein expression, lung inflammation and mortality are reversed when mice are returned to breathing air 1 day following IAV Infection.** Mice were exposed to air or normoxic hypercapnia (10% $CO_2$/21% $O_2$, HC) for 3 days before and continuously after infection with 30 pfu IAV, or to HC for 3 days before and 1 day after IAV infection, followed by air for the remainder of the experiment (A). Viral NS1 protein (red) and nuclei (blue) were visualized by immunofluorescence (IF) microscopy of lung tissue sections from mice sacrificed at 7 dpi (B). Lungs harvested at 4 and 7 dpi were sectioned and stained with H&E (C). Kaplan-Meier plot showing survival after infection (D); N = 6-8 per group, *P<0.05 vs. air by log-rank test.

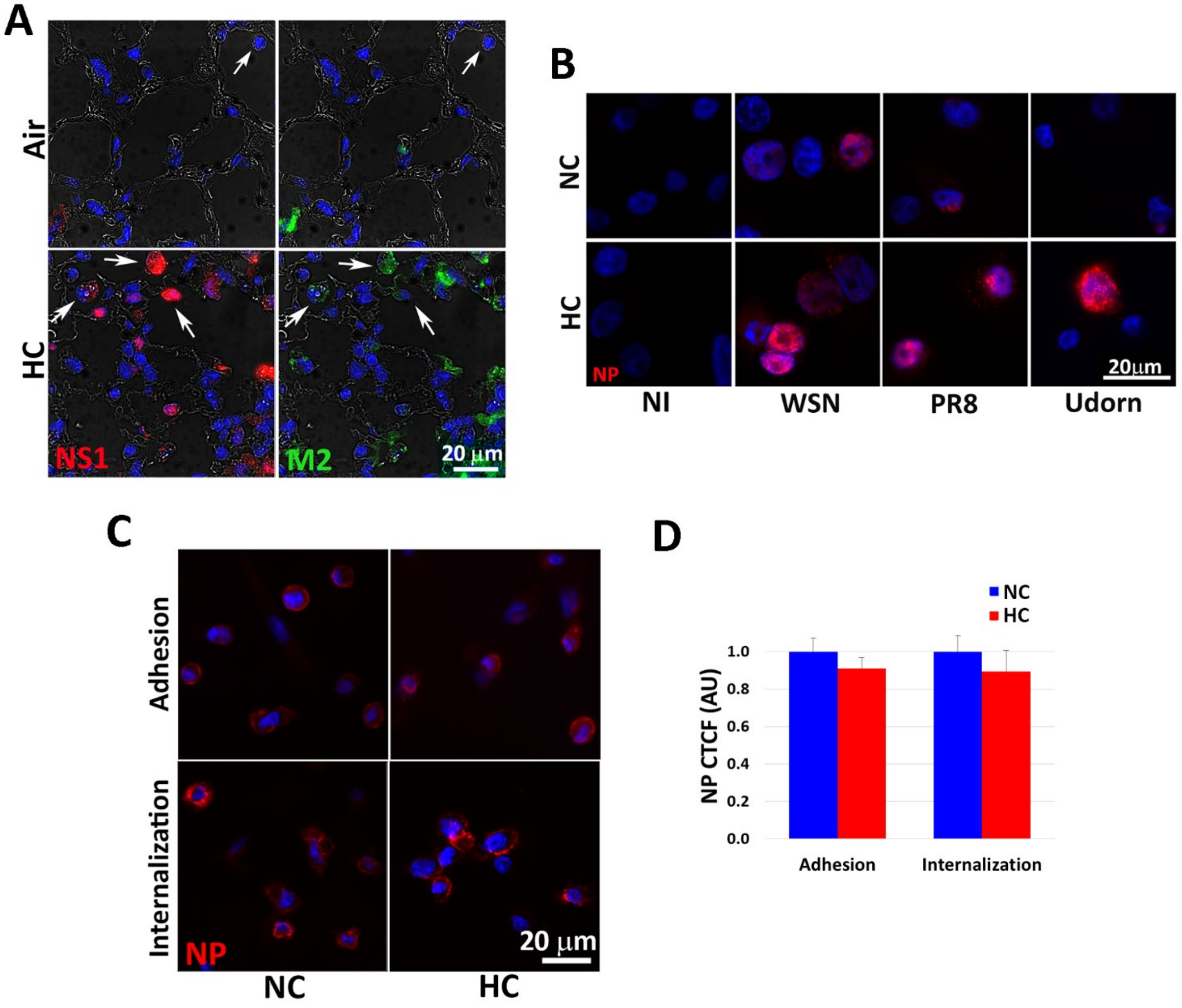


<u>**Fig. S2:**</u> **Hypercapnia increases viral protein expression in lungs from IAV-infected mice, increases NP protein expression in macrophages infected with multiple strains of IAV, and does not change IAV adhesion or internalization in macrophages.** Mice pre-exposed to air or HC and infected with 30 pfu IAV were sacrificed 4 dpi. Viral NS1 (red) and M2 (green) in lung tissue sections were assessed by IF. White arrows indicate AMØs within alveolar airspaces (A). Differentiated THP-1 MØs were cultured in 5% $CO_2$ (NC) or 15% $CO_2$ (HC) for 2 h, infected with A/WSN/1933 [H1N1] (WSN), A/PR8/Puerto Rico/8/1934 [H1N1] (PR8) or A/Udorn/307/1972 [H2N3] (Udorn), at 1, 0.1, and 2 MOI, respectively, or not infected (NI) as control, and cultured in NC or HC for an additional 18 h. Expression of viral NP protein (red) was assessed by IF microscopy (B). Differentiated THP1 MØs in NC or HC for 18 h at 37°C, were then placed on ice, and IAV (20 MOI) was added to the cultures. To assess viral adhesion, after 90 min on ice, cells were washed with ice-cold media and fixed. To assess viral internalization, after 90 min on ice, cells were returned to NC or HC at 37°C for 30 min, then washed and fixed. NP protein (red) was visualized by IF microscopy (C) Nuclei were stained with DAPI (blue). NP was quantified as corrected total cell fluorescence (CTCF), expressed in arbitrary units (AU); N = 3 (D).

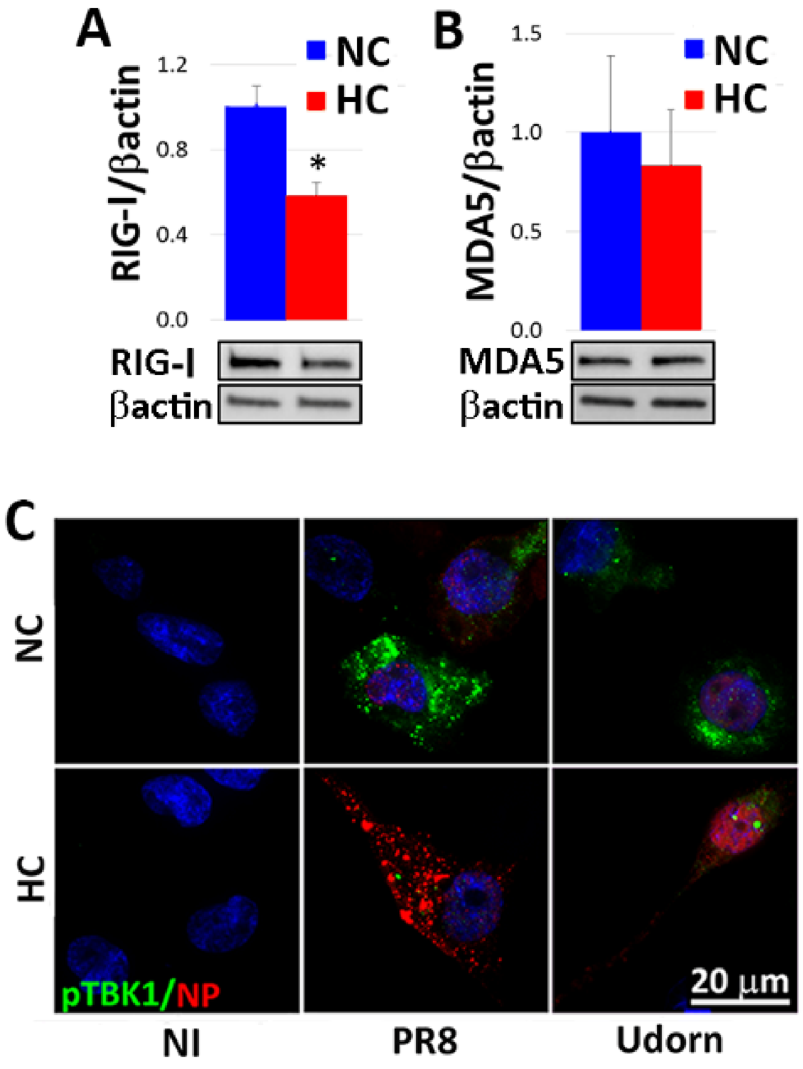


**<u>Fig. S3:</u> Hypercapnia decreases basal RIG1 expression and IAV-induced TBK1 activation.** Differentiated THP-1 MØs were exposed to 5% $CO_2$ (NC) or 15% $CO_2$ (HC) for 18 h and processed for determination of RIG1 (A) and MDA5 (B) by immunoblot; N = 4, *$p < 0.05$. Alternatively, THP-1 MØs were exposed to NC or HC for 2 h, infected with A/PR8/Puerto Rico/8/1934 [H1N1] (PR8) or A/Udorn/307/1972 [H2N3] (Udorn), at 0.1 and 2 MOI respectively, or not infected (NI) as control, and cultured in NC or HC for an additional 18 h. Expression of pTBK1 (green) and viral NP (red) protein was then assessed by IF microscopy; nuclei were stained with DAPI (blue) (C).

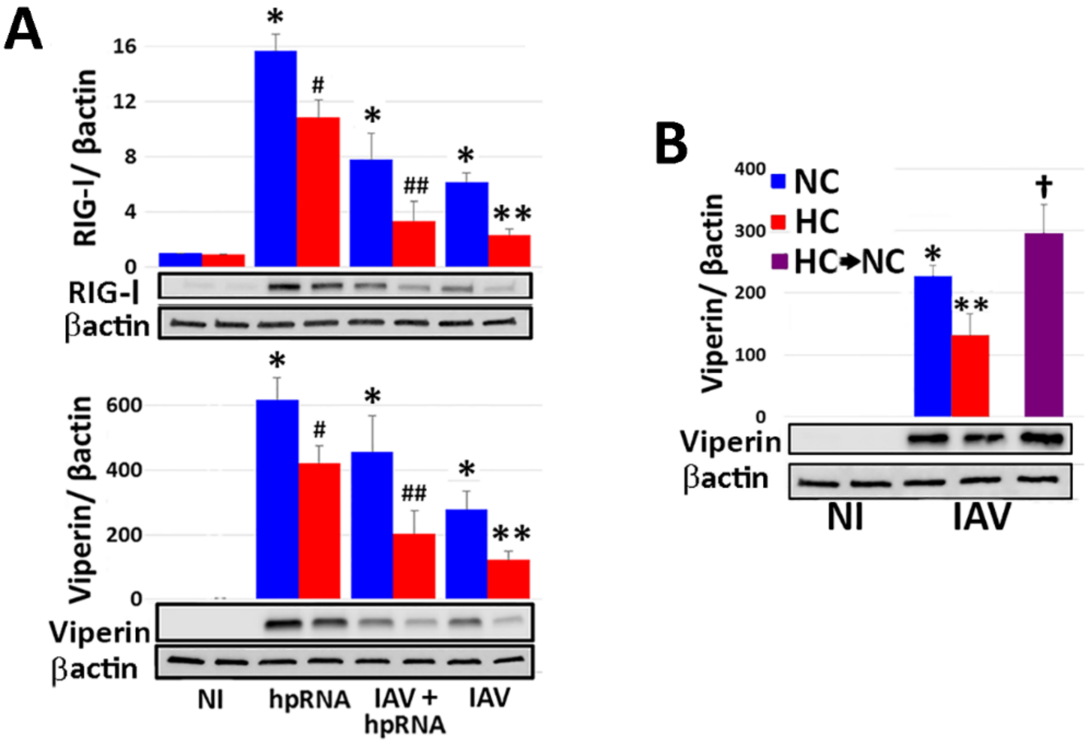


**<u>Fig. S4:</u> Hypercapnia inhibits the macrophage response to the RIG-1 ligand 3p-hpRNA and hypercapnic inhibition of viperin in human MØ is reversible.** Differentiated THP-1 MØs were pre-exposed to 5% $CO_2$ (NC) or 15% $CO_2$ (HC) for 2 h, then stimulated with 3p-hpRNA (hpRNA, 1 µg/ml) and/or infected with IAV (MOI 2), and cultured in NC or HC for an additional 18 h. RIG-1 (top panel) and viperin (bottom panel) protein expression were assessed by immunoblot, with β-actin as loading control. N=5; *P<0.01 vs. NI control, ***p*<0.05 vs. NC + IAV, $^{\#}p<0.05$ vs. NC + hpRNA, $^{\#\#}p<0.05$ vs. HC + hpRNA (A). In addition, THP-1 were cultured continuously in NC, or pre-exposed HC for 18 h, prior to infection with IAV. Cells pre-exposed to HC were then either maintained in HC, or placed in NC (HC → NC), and cultured for an additional 24 h. Viperin was assessed by immunoblot, with β-actin as loading control. N ≥ 3; **p* < 0.01 vs. †NI, ***p* < 0.05 vs. NC + IAV, †*p* < 0.05 vs. HC + IAV (B).

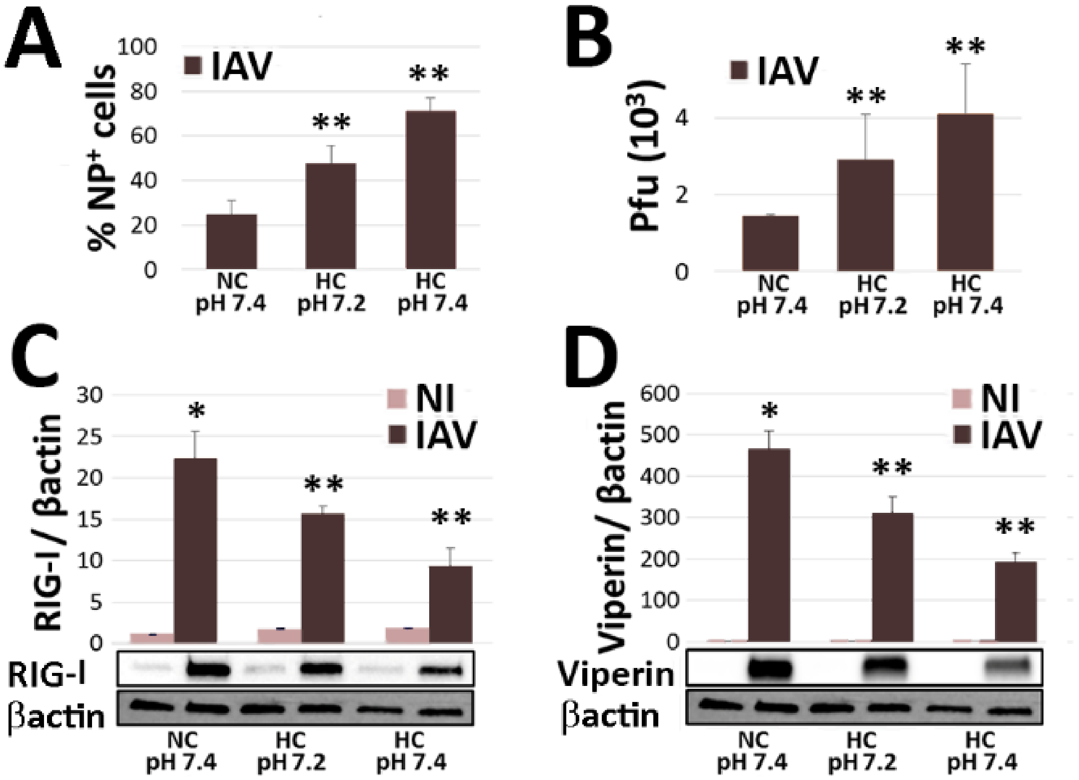


<u>Fig. S5</u>: **The hypercapnia-induced increase in IAV replication and suppression of the IAV-induced antiviral response is not due to extracellular acidosis.** Differentiated THP-1 MØs were pre-exposed to 5% $CO_2$ (NC) or 15% $CO_2$ (HC) for 2 h, infected with IAV, and cultured in NC or HC for an additional 18 h. To assess whether the effects of elevated $CO_2$ were due to extracellular acidosis, cells were exposed to HC in Tris-buffered media at pH 7.2 (equivalent to the pH of 15% $CO_2$-containing unbuffered media) or pH 7.4 (A-D). Viral NP expression was assessed by IF microscopy and quantified as the percentage of NP-positive cells (A). Viral titers in culture supernatants were determined by plaque assay (B). RIG-1 (C) and viperin (D) protein expression were assessed by immunoblot, with β-actin as loading control. N ≥ 3. *$p < 0.01$ vs. NI, **$p < 0.05$ vs. NC + IAV.

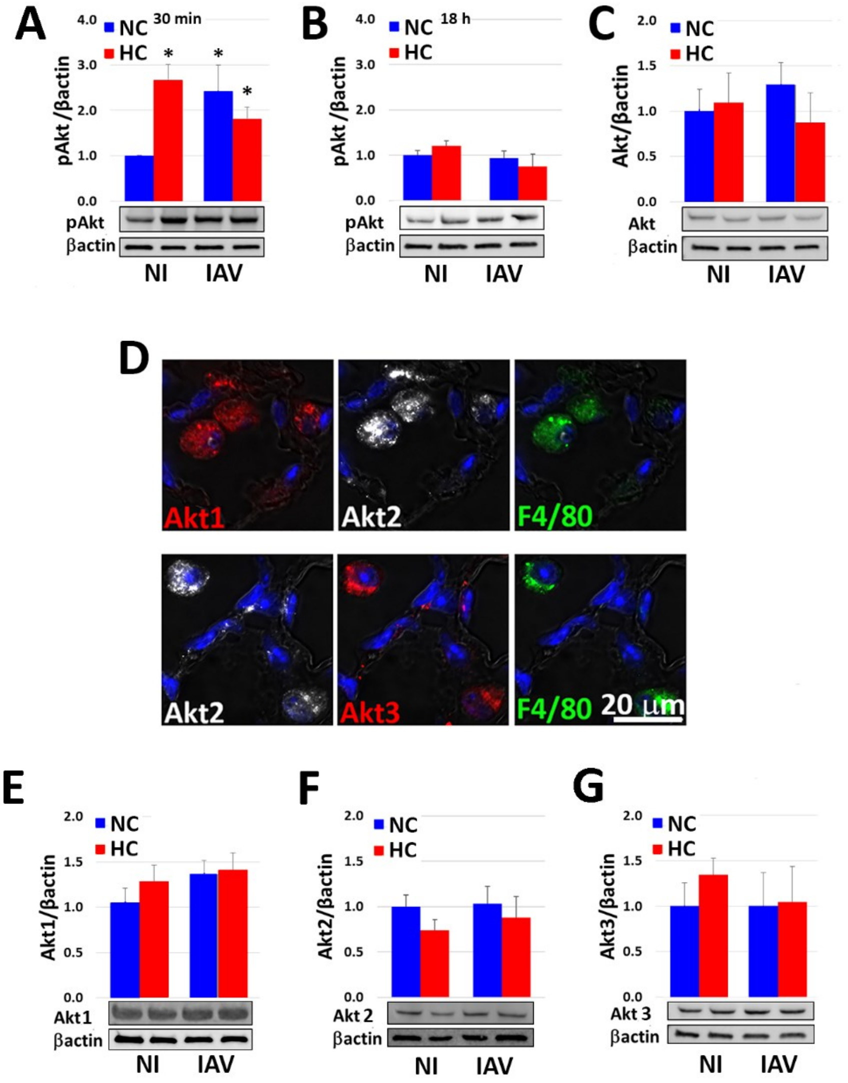


**<u>Fig. S6:</u> Hypercapnia and IAV trigger early activation of Akt at threonine T308/T309/T305 in non-additive fashion but do not alter total Akt, Akt1, Akt2 and Akt3 protein expression.** Differentiated THP-1 MØs were pre-exposed to 5% $CO_2$ (NC) or 15% $CO_2$ (HC) for 2 h, infected with IAV (MOI 2), and cultured in NC or HC for 30 min (A) or 18 h (B) prior to assessment of Akt1/Akt2/Akt3 phosphorylation (pAkt) at threonine T308/T309/T305 by immunoblot. Total Akt was assessed at 18 h by immunoblot using a pan-Akt antibody (C). Immunostaining of lung tissue from untreated mice with isoform-specific antibodies shows that AMØs express Akt1 (red), Akt2 (white) and Akt3 (red); F4/80 (green) was used as MØ marker and nuclei were stained with DAPI (blue) (D). THP-1 MØs cultured in NC or HC for 2 h and infected with IAV (MOI 2) for an additional 18 h prior were assessed of Akt1 (E), Akt2 (F) and Akt3 (G) expression by immunoblot using isoform-specific antibodies; N=4-6.

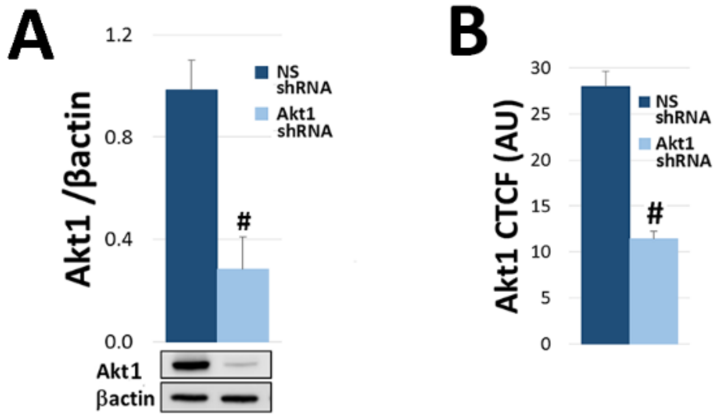


<u>**Fig. S7:**</u> **Akt 1-targeted shRNA decreases Akt1 protein expression in vitro and in vivo.** MLE cells were treated with lentivirus containing non-silencing (NS) shRNA or Akt1-targeted shRNA and cultured for 2 days, at which point expression of Akt1 was determined by immunoblot (A); N = 3, #*p* < 0.05 vs. NS shRNA. Mice were infected intranasally with lentivirus containing NS or Akt1-targeted shRNA, and 14 days later AMØs were collected by BAL and immunostained for Akt1, which was quantified as corrected total cell fluorescence (CTCF), expressed in arbitrary units (AU); N = 3, #*p* < 0.05 vs. NS shRNA (B).

### Supplementary Table 1

| REAGENT or RESOURCE | SOURCE | IDENTIFIER |
|---|---|---|
| **Antibodies** | | |
| Rabbit polyclonal anti-NS1 | GeneTex | Cat#GTX125990; RRID:AB_11170327 |
| Mouse monoclonal anti-IAV M2 | Invitrogen | Cat#MA1-082; RRID:AB_10892134 |
| Mouse monoclonal anti-IAV NP | Abcam | Cat#ab128193; RRID:AB_11143769 |
| Rabbit polyclonal anti-pTBK | Cell Signaling Technology | Cat#5483S; RRID:AB_10693472 |
| Rabbit polyclonal anti-p$^{S473}$Akt | Cell Signaling Technology | Cat#4060S; RRID:AB_2315049 |
| Rabbit monoclonal anti-pAkt$^{Y380}$ | Cell Signaling Technology | Cat#2965S; RRID:AB_2255933 |
| Rabbit monoclonal anti-Akt | Cell Signaling Technology | Cat#4691S; RRID:AB_915783 |
| Mouse monoclonal anti-Akt1 | Cell Signaling Technology | Cat#2967S; RRID:AB_331160 |
| Rabbit polyclonal anti-Akt2 | R&D Systems | Cat#AF23151; RRID:AB_664041 |
| Rabbit polyclonal anti-Akt3 | GeneTex | Cat#GTX113312; RRID:AB_2036200 |
| Rat monoclonal anti-F4/80 | Abcam | Cat#ab6640; RRID:AB_1140040 |
| Goat polyclonal anti-SP-C | Santa Cruz Biotechnology | Cat#Sc-7706; RRID:AB_2185507 |
| Rabbit polyclonal anti-STAT1 | GeneTex | Cat#GTX113010; RRID:AB_10731461 |
| Rabbit polyclonal anti-pSTAT1 | GeneTex | Cat#GTX32266 |
| Rabbit monoclonal anti-RIG1 | Cell Signaling Technology | Cat#3743S; RRID:AB_2269233 |
| Rabbit monoclonal anti-MDA5 | Cell Signaling Technology | Cat#5321S; RRID:AB_10694490 |
| Rabbit monoclonal anti-TRAF3 | Cell Signaling Technology | Cat#61095; RRID:AB_2799601 |
| Rabbit polyclonal anti-Mx1 | Abcam | Cat#ab95926; RRID:AB_10677452 |
| Rabbit polyclonal anti-OAS1 | Abcam | Cat#ab86343; RRID:AB_2158286 |
| Rabbit polyclonal anti-viperin | Abcam | Cat#ab73864; RRID:AB_1640850 |
| Goat anti-Rabbit IgG (H+L) Highly Cross-Adsorbed Secondary Antibody, Alexa Fluor Plus 555 | ThermoFisher | Cat#A32732; RRID:AB_2633281 |
| Donkey Anti-Mouse IgG (H+L) Antibody, Alexa Fluor 488 Conjugated | ThermoFisher | Cat#A21202; RRID:AB_141607 |
| Rabbit anti-Goat IgG (H+L) Superclonal(TM) Secondary Antibody, Alexa Fluor 647 | ThermoFisher | Cat#A27018; RRID:AB_2536082 |
| LI-COR IRDye 680RD Goat anti-Mouse | ThermoFisher | Cat#NC0252290 |
| Anti-rabbit IgG, HRP-linked Antibody | Cell Signaling Technology | Cat#7074; RRID:AB_2099233 |
| Anti-mouse IgG, HRP-linked Antibody | Cell Signaling Technology | Cat#7076; RRID:AB_330924 |

| | | |
|---|---|---|
| **Bacterial and Virus Strains** | | |
| Mouse-adapted influenza A virus (A/WSN/33 [H1N1]) | Robert Lamb Lab | |
| IAV WSN/1933(H1N1) | Karen Ridge Lab | |
| Udorn/307/1972(H2N3) | | |
| PR8/Puerto Rico/8/1934 (H1N1) | ATCC | Cat#VR-1469 |
| VSVG pseudotyped lentiviruses | Gene Hunter Corporation | |
| **Chemicals, Peptides, and Recombinant Proteins** | | |
| Recombinant Mouse IFN-beta Protein | R&D System | Cat#8234-MB-010 |
| MK2206 | MedChemExpress LLC | Cat#HY-10358 |
| Captisol | MedChemExpress LLC | Cat#HY-17031 |
| A-674563 (2mg)-AKt1 inhibitor | Selleckchem | Cat#S2670 |
| CCT128930 (5mg)--Akt2 inhibitor | Selleckchem | Cat#S2635 |
| LY294002 | Selleckchem | Cat#S1105 |
| Recombinant Mouse IFN- Gamma (Animal-Free) | ThermoFisher | Cat#50-712-661 |
| **Experimental Models: Cell Lines** | | |
| MDCK cells | ATCC | Cat#ATCC® CCL-34 |
| Mouse bronchoalveolar lavage (BAL) AMØs | In this paper | |
| Human AMØs | Northwestern University | |
| Human monocytic leukemia THP-1 cells | ATCC | Cat#ATCC® TIB-202 |
| MLE cells | ATCC | Cat#ATCC® CRL-2110 |
| 293T packaging cells | Gene Hunter Corporation | |
| **Experimental Models: Organisms/Strains** | | |
| C57Bl/6 mouse | Jackson Laboratory | Cat#C57Bl/6 |
| **Oligonucleotides** | | |
| RIG1 (Hs00204833_m1) | Thermo Fisher | Cat#4331182 |
| IFNβ (Hs01077958_s1) | Thermo Fisher | Cat#4331182 |
| Mx1 (Hs00895608_m1) | Thermo Fisher | Cat#4331182 |
| MDA5 (Hs.PT.58.1224165) | IDT | NA |
| IFNAR1 (Hs.PT.58.20048943) | IDT | NA |
| OAS1 (Hs.PT.58.2338899) | IDT | NA |
| viperin (Hs.PT.58.713843) | IDT | NA |
| IFNAR2 (Hs.PT.58.1621113) | IDT | NA |
| SREBF2 (Hs.PT.58.45335433) | IDT | |
| ABCA1 (Hs.PT.58.27452429) | IDT | |
| HMGCR (Hs.PT.58.41105492) | IDT | |
| GAPDH (Hs99999905_m1) | Thermo Fisher | Cat#4331182 |
| **Recombinant DNA** | | |
| Packaging vectors psPAX2 | Addgene | |
| Packaging vectors pMD2.G | Addgene | |
| Third generation lentiviral expression vector pLKO | Sigma-Aldrich | |
| shRNAs clones against mouse Akt1 | Sigma-Aldrich | Cat# TRCN0000304735, TRCN0000304683, TRCN0000304684, TRCN0000302357, TRCN0000304736 |
| Non-silencing control shRNA | Sigma-Aldrich | Cat#SHC002 |
